# Noninvasive Injectable Optical Nanosensor-Hydrogel Hybrids Detect Doxorubicin in Living Mice

**DOI:** 10.1101/2023.12.05.570137

**Authors:** Zachary Cohen, Dave J. Alpert, Adam C. Weisel, Arantxa Roach, Syeda Rahman, Pooja V. Gaikwad, Steven B. Nicoll, Ryan M. Williams

## Abstract

While the tissue-transparent fluorescence of single-walled carbon nanotubes (SWCNTs) imparts substantial potential for use in non-invasive biosensors, development of non-invasive systems is yet to be realized. Here, we investigated the functionality of a SWCNT-based nanosensor in several injectable SWCNT-hydrogel systems, ultimately finding SWCNT encapsulation in a sulfonated methylcellulose hydrogel optimal for detection of ions, small molecules, and proteins. We found that the hydrogel system and nanosensor signal were stable for several weeks in live mice. We then found that this system successfully detects local injections of the chemotherapeutic agent doxorubicin in mice. We anticipate future studies to adapt this device for detection of other analytes in animals and, ultimately, patients.

## Introduction

Single-walled carbon nanotubes (SWCNTs), cylindrical allotropes of carbon, have garnered considerable research interest in recent years for their exceptional mechanical, electrical, and optical properties.^1^ In particular, semiconducting SWCNTs exhibit near-infrared fluorescence, making them well-suited to use as transducers in optical biosensors. SWCNTs exhibit large stokes shifts,^2^ do not photobleach,^3, 4^ and fluoresce in the tissue-transparent window. This has enabled implanted SWCNT-based biosensors to retain their fluorescence properties for at least 300 days in vivo,^5^ and at depths of 5.5 cm ex vivo.^6^ SWCNT-based biosensors have been developed for a variety of analytes, including metals,^7-9^ nitrous oxide,^10^ glucose,^11^ chemotherapeutics,^12, 13^ auxins,^14^ insulin^15^ and other hormones,^16, 17^ neurotransmitters,^18-20^ oligonucleotides,^21^ riboflavin,^17^ fibrinogen,^22^ growth factors,^23^ amyloid beta,^24^ cytokines,^25^ lipid accumulation diseases,^26^ cardiac biomarkers,^27^ cancer biomarkers,^28, 29^ and more.^30^

To date, several studies have demonstrated SWCNT-based biosensor implants in living animals. SWCNTs can be administered intravenously,^31^ which results in SWCNT uptake in the liver Kupffer cells, allowing sensing within those cells.^32^ For biosensing in other locations, the majority of studies surgically implanted SWCNTs in either sealed dialysis membranes^12, 21, 28^ or hydrogels^5, 6, 13, 16, 33-35^ under anesthesia. Surgical implantation makes biosensing possible at any location, but is by nature an invasive procedure. However, an injectable SWCNT implant, in which SWCNT are encapsulated in an in situ-gelling hydrogel, would enable noninvasive biosensing without necessitating an invasive procedure. One prior study has taken a similar approach in marine organisms, delivering nanotubes dispersed in pre-gelled polymers via trocar.^36^ Though innovative, this approach requires larger trochar needles for pre-formed gels and may be better adapted to the clinic by instead using polymers which gelate in situ post-injection, thereby conforming to nearby tissue. Such injectable hydrogels are already used for cell and drug delivery.^37^

Hydrogels are a diverse and customizable class of materials. Their chemical properties may be tuned to control their mechanical properties, porosity, biocompatibility, and other properties relevant to injection at numerous physiological locations.^37^ They have such diverse application as platforms for drug and cell delivery, cell recruitment, and as scaffolds for tissue engineering. Their high degree of tunability is important for SWCNT implants in particular, as such a device must facilitate analyte diffusion into the gel while preventing SWCNT diffusion out of the gel.

Other studies have demonstrated impressive performance for surgically implanted hydrogel-encapsulated SWCNTs. Beyond preserving SWCNT fluorescence for 300 days,^5^ multiple experiments have shown that hydrogels can reliably sequester SWCNTs without leakage.^35, 38^ Furthermore, hydrogel-encapsulated SWCNTs have been used to detect such diverse analytes as nitrous oxide,^5, 33^ progesterone,^16, 35^ riboflavin,^34, 35^ ascorbate,^34^ and chemotherapeutics^13^ in vivo. These studies commonly employed gels incorporating alginate^5, 33^ or poly(ethylene glycol) diacrylate,^13, 16, 34, 35^ polymers noted for their biocompatibility.

In this work, we developed a novel injectable hydrogel-encapsulated SWCNT system based on methacrylated methylcellulose (MC).^39^ We compared this system to previously-described alginate and chitosan encapsulation systems ^40,41^. We found that this novel injectable system responds robustly to a variety of analytes, including magnesium chloride, sodium bicarbonate, the chemotherapeutic doxorubicin (DOX), and bovine serum albumin. We then found that MC-SWCNTs implanted via simple, noninvasive injection show stable fluorescence for several weeks. Finally, we found that the sensor system responds to DOX in living mice as a model of a clinically-relevant analyte.

## Methods

### Sensor Preparation

(GT)_15_-SWCNTs were prepared as previously described.^28, 42^ Briefly, in a 1.5 mL microcentrifuge tube, approximately 0.5 mg HiPCO SWCNTs (NanoIntegris, Boisbriand, Quebec, Canada) and (GT)_15_ single-stranded DNA (10 g/L, Integrated DNA Technologies, Coralvile, IA, USA) suspended in 1X PBS (phosphate-buffered saline, Sigma-Aldrich, St. Louis, MO, USA) were added to a final oligonucleotide:SWCNT mass ratio of 2:1. 1X PBS was added to bring the final volume to 500 µL. To achieve aqueous dispersion of SWCNTs by (GT)_15_, the solution was sonicated (1 hr., 40% amplitude) in an ice bath by a VCX 750 (Sonics & Materials, Inc., Newtown, CT, USA) fitted with a 2 mm stepped-microtip. The suspension was then ultracentrifuged (58,000 × g, 1 hr., 4 °C) in an Optima MAX-XP (Beckman Coulter, Indianapolis, IN, USA) to separate residual catalyst, amorphous carbon, and any partially suspended SWCNTs from the (GT)_15_-SWCNTs. The top 75% of the solution was stored at 4 °C until further use.

Prior to an experiment, excess (GT)_15_ was removed by filtration. An aliquot (100-375 µL) of the (GT)_15_-SWCNT solution was loaded into a 100 kDa molecular weight cutoff filter (Millipore Sigma, St. Louis, MO, USA) and centrifuged (14,000 RPM, 15 min, 4 °C) in a Sorvall ST8R centrifuge (Thermo Fisher Scientific, Waltham, MA, USA). The filtrate was discarded, the contents of the filter were resuspended in 1X PBS (500 µL), and centrifugally filtered again. (GT)_15_-SWCNTs remaining in the filter were resuspended in 1X PBS (100-200 µL).

The resultant solution was characterized by absorbance spectroscopy as previously described.^28, 42^ Briefly, a visible absorbance spectrum was acquired from the (GT)_15_-SWCNTs using a V-730 UV-VIS spectrophotometer (JASCO, Easton, MD, USA). Concentration was determined using an empirically derived extinction coefficient of 0.036 L mg^-1^ cm^-1^ for the absorption minimum at ~ 630 nm.^27^

### Fluorescence Spectrum Acquisition & Analysis

Final (GT)_15_-SWCNT concentrations were 1 mg/L across all experiments. For experiments using the NS MiniTracer (Applied NanoFluorescence, TX, USA), samples were excited using a 50 mW 638 nm laser with 1-3 s exposure time. For experiments using the ClaIR IR plate reader (Photon Etc., Montreal, Quebec, Canada), samples were excited at 655 and 730 nm (sequentially) for 0.5 s at 1.75 W. For in vivo experiments using the IRina in vivo NIR II spectral probe (Photon Etc.), spectra were acquired with de-focused 1 W laser excitation at 655 and 730 nm (sequentially), 1 s exposures, 1 × 2 binning, manual dark subtraction, and manual autofluorescence subtraction. Autofluorescence was modeled as exponential decay.

Fluorescence peaks were assigned to chiralities based on optical characterization in the literature.^43^ To determine the center wavelength and intensity of the (7,5) E_11_ peak, the 24 points closest to the emission maximum around 1035-1045 nm were fit to a pseudo-Voigt model (Figure S1, Equation S1) using custom MATLAB code. Fits of other chiralities were performed similarly, using consistent numbers of points for each peak (14 points for less prominent peaks, 28 points for broader peaks). Fits of binned spectra were performed using half as many data points. All analyses presented used fits deemed to be of sufficiently high quality (R^2^ ≥ 0.9).

### Magnesium Chloride (MgCl_2_) and Sodium Bicarbonate (NaHCO_3_) Detection in Alginate-Acrylamide Gels

(GT)_15_-SWCNT-containing (1.0 mg/L) alginate-acrylamide gels were prepared by modifying a previously published procedure.^40^ Sodium alginate (116.7 mg, Thermo Fisher Scientific) and n-isopropylacrylamide (1524.94 mg, Thermo Fisher Scientific) were added to a glass vial containing (GT)_15_-SWCNTs in 1X PBS (1 mg/L, 20 mL). The vial’s contents were dissolved by slow magnetic stirring until visibly homogenous. Ascorbic acid (40 mg, Sigma-Aldrich) was added to the vial, which was then vortexed. Ammonium persulfate (50 mg, Thermo Fisher Scientific) was added to the vial, which was vortexed again and allowed to sit at room temperature for 15 minutes before being transferred to storage (4 °C).

1 mL of the resultant solution was transferred to 18 cuvettes, which were covered with parafilm and incubated in a water bath (0.5 hr., 40 °C) to gel the alginate-acrylamide copolymer. Fluorescence spectra were acquired from each sample using an NS MiniTracer (1 s exposure). Samples were then divided into three groups (n = 6) and topped with 1 mL of either 1X PBS, magnesium chloride (MgCl_2_, 5 M, Thermo Fisher Scientific), or sodium bicarbonate (NaHCO_3_, 1 M, Thermo Fisher Scientific). Fluorescence spectra were again acquired using an NS MiniTracer (1 s exposure) after one hour of covered incubation in the water bath.

### MgCl_2_ and NaHCO_3_ Detection in Chitosan Gels

(GT)_15_-SWCNT-containing (1.5 mg/L) chitosan gels were prepared by modifying a previously published procedure.^41^ Deionized water (5955.5 µL) and acetic acid (4.5 µL, Sigma-Aldrich) were added to a glass vial containing chitosan (200 mg, Sigma-Aldrich). The resulting mixture was dissolved by magnetic stirring at 40 °C for at least an hour. To this mixture we added glycerol 2-phosphate (3 M, Thermo Fisher Scientific) and (GT)_15_-SWCNTs (3.75 mg/L) in deionized water (4 mL). The resultant solution was slowly mixed magnetically for at least an hour.

1 mL of the resultant solution was transferred to each of nine cuvettes. The samples were covered with parafilm and gelled by incubation in a water bath (40 °C, 1 hr). Fluorescence spectra were acquired using an NS MiniTracer (3 s exposure). The samples were then split into three groups and topped with 1 mL of deionized water, MgCl_2_ (5 M), or NaHCO_3_ (1 M). Fluorescence spectra were again acquired after one hour of covered incubation in the water bath. This procedure was repeated for a total of six samples per group.

### Methylcellulose-SWCNT Gel Preparation

(GT)_15_-SWCNT-contianing MC gels were prepared by modifying our previously published procedure.^39^ Briefly, methylcellulose (Sigma-Aldrich) was methacrylated via the esterification of monomeric hydroxyl groups with methacrylic anhydride (Sigma-Aldrich) and then lyophilized for storage. Methacrylated MC was then dissolved in 1X Dulbecco’s phosphate buffered saline (Thermo Fisher Scientific) containing (GT)_15_-SWCNTs (1 mg/L). For sulfonated gels, 2-sulfoethyl methacrylate (PolySciences, Warminster, PA) was added to a final concentration of 5 mM. The resultant polymer-nanotube solution was then split in half to add redox initiators; to one half was added ammonium persulfate (final concentration 10 mM) and to the other was added an equimolar amount of ascorbic acid (Sigma-Aldrich). These solutions were then transferred to a dual barrel syringe fitted with a mixing tip (such that each barrel contained polymers, nanotubes, and one of the redox initiators). Gels were cast by extrusion through the mixing tip.

### MgCl_2_ Detection in Methylcellulose Gels

2% and 3% MC gels containing (GT)_15_-SWCNTs (1.0 mg/L) were cast into pre-weighed cuvettes (250-500 mg gel per cuvette). After 30 minutes, cuvettes were weighed again and gels were topped with 1X PBS as control or MgCl_2_ (5 M) in 1X PBS and then covered with parafilm. Solution volumes were 1 µL per mg gel. Fluorescence spectra were acquired using an NS MiniTracer (3 s exposure) immediately, every hour for the next three hours, and every day for the next three days. Samples were stored at 37 °C between measurements.

### NaHCO_3_ Detection in Methylcellulose Gels

2% and 3% MC gels containing (GT)_15_-SWCNTs (1.0 mg/L) were cast into cuvettes (250-500 mg gel per cuvette). After 30 minutes of incubation, gels were topped with 1X PBS (1 mL) as control or with NaHCO_3_ (1 M, 1 mL) and then covered with parafilm (n = 4). Fluorescence spectra were acquired with an NS MiniTracer (1 s exposure) immediately upon topping, after one hour, and after two days. Samples were stored at 37 °C between measurements.

### MgCl_2_ Quantification in MC Gels

2% and 3% MC gels containing (GT)_15_-SWCNTs (1.0 mg/L) were cast into pre-weighed cuvettes (250-500 mg gel per cuvette, 20 cuvettes total). After 30 minutes, cuvettes were weighed. For each MC concentration group (n = 5 – 6), one cuvette was topped with 1X PBS as a control, and the nine remaining cuvettes were topped with MgCl_2_ solutions in 1X PBS whose concentrations ranged from 1 M to 3.9 mM, decreasing by half. Solution volumes were 1 µL per mg gel. Cuvettes were covered with parafilm and stored at 37 °C between measurements. Fluorescence spectra were acquired with an NS MiniTracer (3 s exposure) 1, 2, and 6 days after gels were topped.

### MgCl_2_ and Bovine Serum Albumin Detection in Sulfonated Methylcellulose Gels

2% and 3% MC and MC-SO_3_ gels containing (GT)15-SWCNTs (1 mg/L) were cast into 54 pre-weighed cuvettes (250-500 mg gel per cuvette). After 30 minutes, cuvettes were weighed, and topped with 1X PBS as control, MgCl_2_ (5 M), or bovine serum albumin (BSA, 600 µM) and then covered with parafilm (n = 5 – 6). Solution volumes were 1 µL per mg gel. Fluorescence spectra were acquired using an NS MiniTracer (10 s exposure) 1-, 2-, and 6-days post-topping. Samples were stored at 37 °C between measurements. Separately, a set of identical experiments were performed to monitor BSA detection up to two weeks (n = 7) with spectra acquired at 1, 2, 3, 4, 7, 8, and 14 days.

### BSA Quantification in Sulfonated Methylcellulose Gels

3% MC-SO_3_ gels containing (GT)_15_-SWCNTs (1 mg/L) were cast into a 96-well plate (150 ± 50 µL per well, 24 wells). After 30 minutes, fluorescence spectra were acquired using a ClaIR IR plate-reader, then these wells were topped with solutions of BSA in 1X PBS (600, 120, 24, 4.8, 0.96, 0.192, 0.0384, and 0 µM; 150 µL; 3 wells per concentration). Fluorescence spectra were acquired immediately after topping, 1, 2, 3, 4, 5, 24, and 48 hours thereafter.

Solutions of (GT)_15_-SWCNTs in 1X PBS (2 mg/L) were added to other wells in the same plate (150 µL per well, 21 wells). Fluorescence spectra were acquired before the addition of BSA solutions in 1X PBS (240, 48, 9.6, 1.92, 0.384, .0768, and 0 µM; 150 µL; 3 wells per concentration). Another set of wells was filled with (GT)_15_-SWCNTs (1 mg/L) and BSA (600 µM) in 1X PBS (300 µL, 3 wells). Fluorescence spectra were acquired immediately, 1, 2, 3, 4, 5, 24, and 48 hours thereafter.

### Doxorubicin Quantification in Sulfonated Methylcellulose Gels

3% MC-SO_3_ gels containing (GT)_15_-SWCNTs (1 mg/L) were cast into a 96-well plate (150 ± 50 µL per well, 24 wells). After 30 minutes, fluorescence spectra were acquired using a ClaIR IR plate-reader, then these wells were topped with solutions of doxorubicin (DOX) in 1X PBS (1000, 200, 40, 8, 1.6, 0.32, 0.064, and 0 µM; 150 µL; 3 wells per concentration). Fluorescence spectra were acquired immediately after topping, 1, 2, 24, and 48 hours thereafter.

Solutions of (GT)_15_-SWCNTs in 1X PBS (2 mg/L) were added to other wells in the same plate (150 µL per well, 21 wells). Fluorescence spectra were acquired before the addition of DOX solutions in 1X PBS (200, 40, 8, 1.6, 0.32, 0.064, and 0 µM; 150 µL; 3 wells per concentration). Another set of wells was filled with (GT)_15_-SWCNTs (1 mg/L) and DOX (1000 µM) in 1X PBS (300 µL, 3 wells). Fluorescence spectra were acquired immediately, 1, 2, 24, and 48 hours thereafter.

### Animal Studies

Studies in animals were performed in 4-6 week old female SKH1-Elite mice (Crl:SKH1-*Hr*^*hr*^; Charles River, Wilmington, MA). Animals were housed under standard light/dark (12/12) conditions with *ad libitum* access to food and water. All experiments were approved by the Institutional Animal Care and Use Committee of The City College of New York.

### SWCNT-Gel Fluorescence Stability In Vivo

To evaluate the suitability of 3% MC-SO_3_ as a SWCNT injection platform, a mouse was injected subcutaneously in the dorsal region with sterile 3% MC-SO_3_ containing (GT)_15_-SWCNTs. After 30 minutes, the mouse was anesthetized with isoflurane and spectra were acquired using an IRina NIR II spectral probe. Spectra were again acquired in this fashion after 3, 7, 11, and 61 days.

### DOX Detection In Vivo

To evaluate the response of (GT)_15_-SWCNTs in 3% MC-SO_3_ to DOX in vivo, ten mice were dorsally injected with the sensor-containing gel using a dual barrel syringe for subcutaneous injection. Fluorescence spectra were acquired using an IRina in vivo NIR II spectral probe from each mouse under anesthesia 30 minutes post-injection, after which they were immediately subcutaneously dosed with 10 mg/kg DOX in 1 mL 1X PBS or 1 mL 1X PBS as control (n = 3). As mice masses ranged from 18-24 g, doses ranged from 330 to 440 nmol (330 - 440 µM). Doses were administered in quarters at four locations spaced around the implant site (4 × 250 µL). Spectra were acquired immediately, 10 minutes, 4 hours, 24 hours, and 48 hours after dosing.

### Statistical Analyses

Statistical analysis to determine sensor response significance was performed via two-tailed t-tests with unequal variances.

## Results and Discussion

The fluorescence of SWCNT-based nanosensors was evaluated in six hydrogels. SWCNT fluorescence was detected in all gels tested (Figure 1). As expected, we found that nanotube fluorescence in relatively hydrophilic gels (e.g., chitosan) was red-shifted relative to those in relatively hydrophobic gels (e.g., methylcellulose).^44, 45^

**Figure 1.**
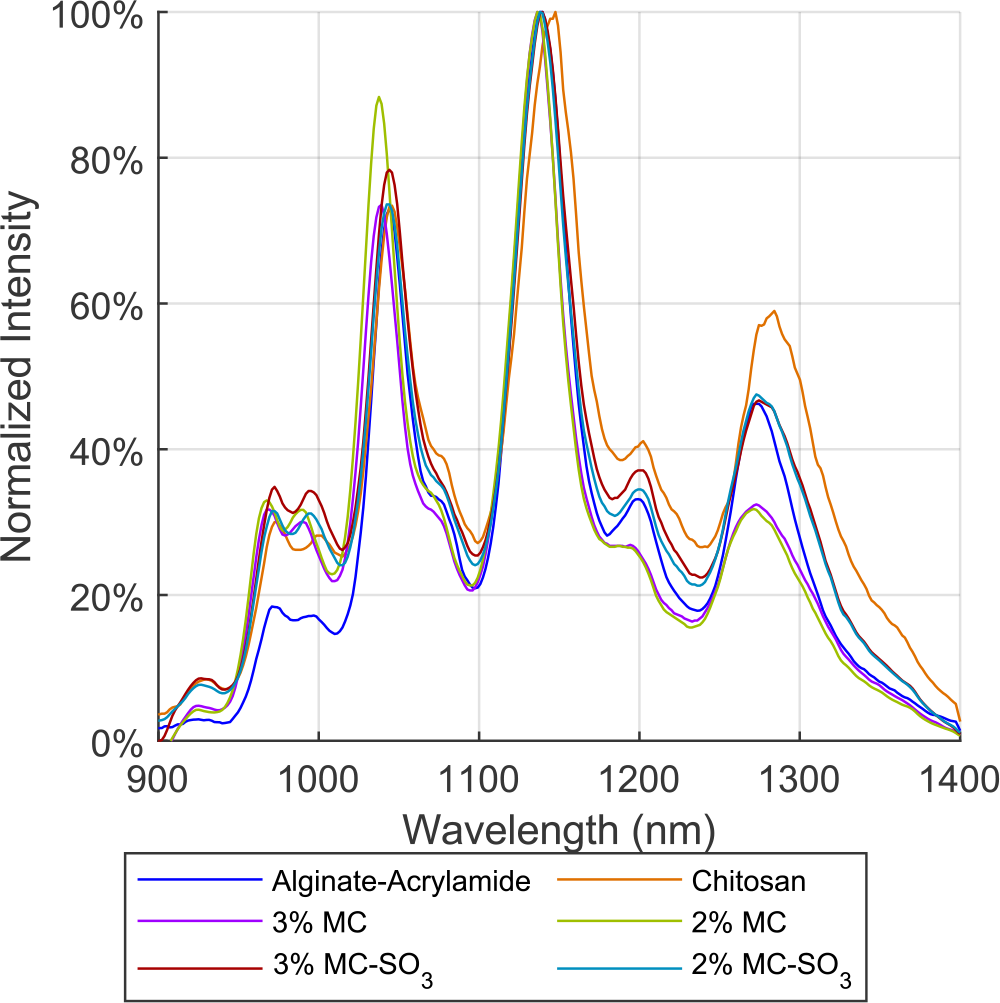
Representative Fluorescence Spectra of the Nanotube Sensor in All Hydrogels Tested.

## Detection of Magnesium Chloride and Sodium Bicarbonate in Alginate-Acrylamide Gels

The first hydrogel tested was a previously described alginate-acrylamide copolymer solution that crosslinks at body temperature.^40^ The copolymer is composed primarily of poly(*N*-isopropylacrylamide) grafted to an alginate backbone for extra rigidity.. It was selected for these experiments because alginate gels are commonly used in experiments with SWCNTs, while crosslinking at body temperature is appealing for injection.^5, 12, 33, 37^ As a naturally-occurring polysaccharide, alginate is highly abundant and biocompatible.^46^ It is also known for forming gels relatively easily, especially via the addition of a divalent crosslinker.

The sensing ability of (GT)_15_-SWCNTs in the alginate-acrylamide gel was evaluated using magnesium chloride (MgCl_2_, 5 M) and sodium bicarbonate (NaHCO_3_, 1 M) as representative analytes. MgCl_2_ and NaHCO_3_ were chosen for this initial response evaluation because their small hydrodynamic radii and high charge were hypothesized to better facilitate diffusion and detection. MgCl_2_ was of particular interest as previous studies have demonstrated good detection of divalent cations by SWCNTs.^7-9^ After SWCNT-containing alginate-acrylamide gels were incubated with the analytes for one hour at body temperature, NaHCO_3_ was found to cause partial quenching (intensity decreased by 53 ± 54% versus control) and blue-shifting (-2.3 ± 1.8 nm). MgCl_2_ only caused shifting (2.3 ± 0.6 nm, Figure 2), though this shift was more consistent than that induced by NaHCO_3_.

**Figure 2.**
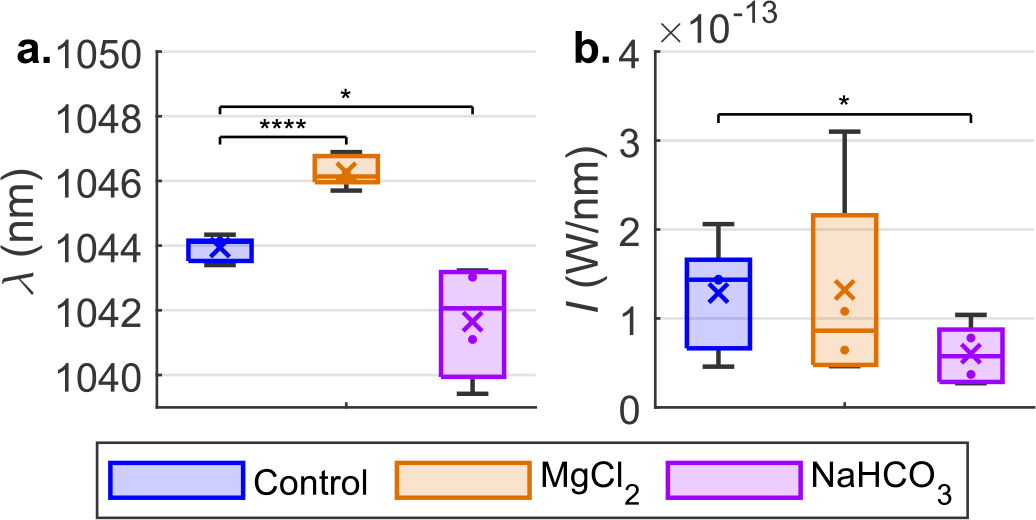
Sensor Response to MgCl_2_ and NaHCO_3_ in alginate-acrylamide hydrogels. A) center wavelength and b) intensity. MgCl_2_ and NaHCO_3_ induced significant wavelength shifting versus control (p = 4 × 10^-6^ and .02 respectively), while only NaHCO_3_ caused a significant intensity change (p = 0.04). N = 6.

### Detection of MgCl_2_ and NaHCO_3_ in Chitosan Gels

We then investigated sensor encapsulation in a chitosan gel, also known to crosslink at body temperature, whose mechanical properties were previously shown to improve upon incorporation of graphene oxide.^41^ Similar to alginate, chitosan was an appealing polymer because of its abundance and biocompatibility.^46^ The sensing ability of (GT)_15_-SWCNTs in the chitosan gel was again evaluated using MgCl_2_ (5 M) and NaHCO_3_ (1 M) as model analytes (Figure 3). While neither caused an intensity-based response, MgCl_2_ induced red-shifting (6.1 ± 1.1 nm) which plateaued after an hour and then decreased by a third (to 4.2 ± 1.4) nm after the first day. We found minimal response to NaHCO_3_, likely because the gel is itself basic.

**Figure 3.**
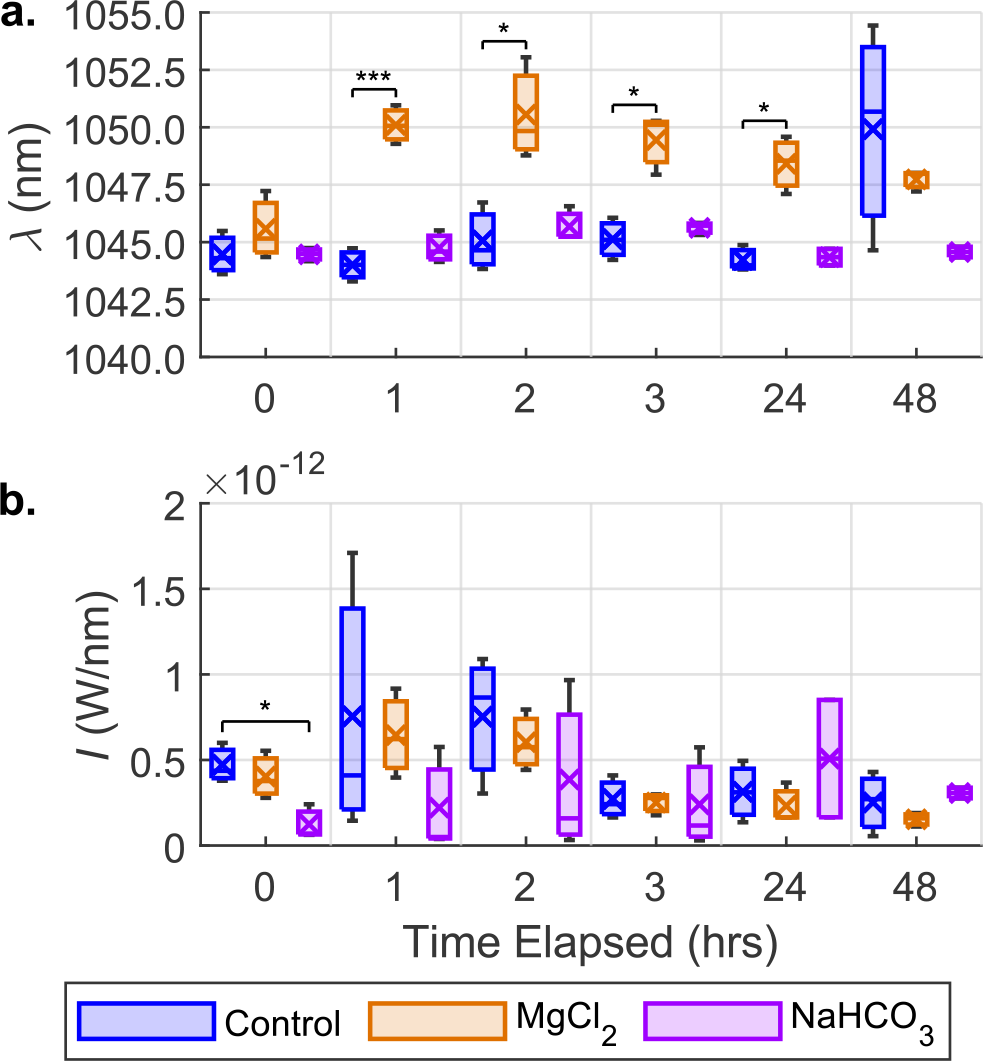
Sensor Response to MgCl_2_ and NaHCO_3_ in a chitosan-based hydrogel. A) center wavelength and b) intensity. NaHCO_3_ did not cause shifting at any timepoint; MgCl_2_ induced significant shifting on hours 1, 2, 3, and 24 (p = 8 × 10^-4^, .03, .01, and .02 respectively). No significant differences were observed in intensities. N = 3.

### Detection of MgCl_2_ and NaHCO_3_ in Methylcellulose Gels

While both the chitosan and alginate-acrylamide gels gave acceptable results for model analyte detection, they are known to be less mechanically robust than desirable for long-term implantation.^40, 41^ This prompted us to investigate a gel based on methacrylated methylcellulose (MC) with demonstrated biocompatibility and mechanical properties facilitating long-term in vivo stability.^39^ This particular gel was originally investigated for tissue engineering applications, and therefore is already well-suited to clinical translation. Additionally, cellulose is the most naturally abundant polysaccharide and is therefore easy to source.^46^

MC gels were initially tested at polymer concentrations of 2% and 3%. While 3% MC gels were previously shown to have optimal mechanical properties for implantation as an adipose tissue mimic, 2% MC gels were speculated to better facilitate analyte diffusion because of their higher porosity.^39^ Though neither MC-SWCNT formulation responded to MgCl_2_ (5 M) by the first day, large red shifts were observed two days into the experiment (Figure 4, 7.5 ± 0.2 nm and 6.6 ± 0.2 nm for 3% and 2% gels respectively). While 2% MC was not found to perform better than 3% MC for MgCl_2_ detection, a similar experiment found that NaHCO_3_ (1 M) only elicited a response from the 2% system, inducing red shifting two days after exposure (Figure 5, 1.6 ± 0.5 nm).

**Figure 4.**
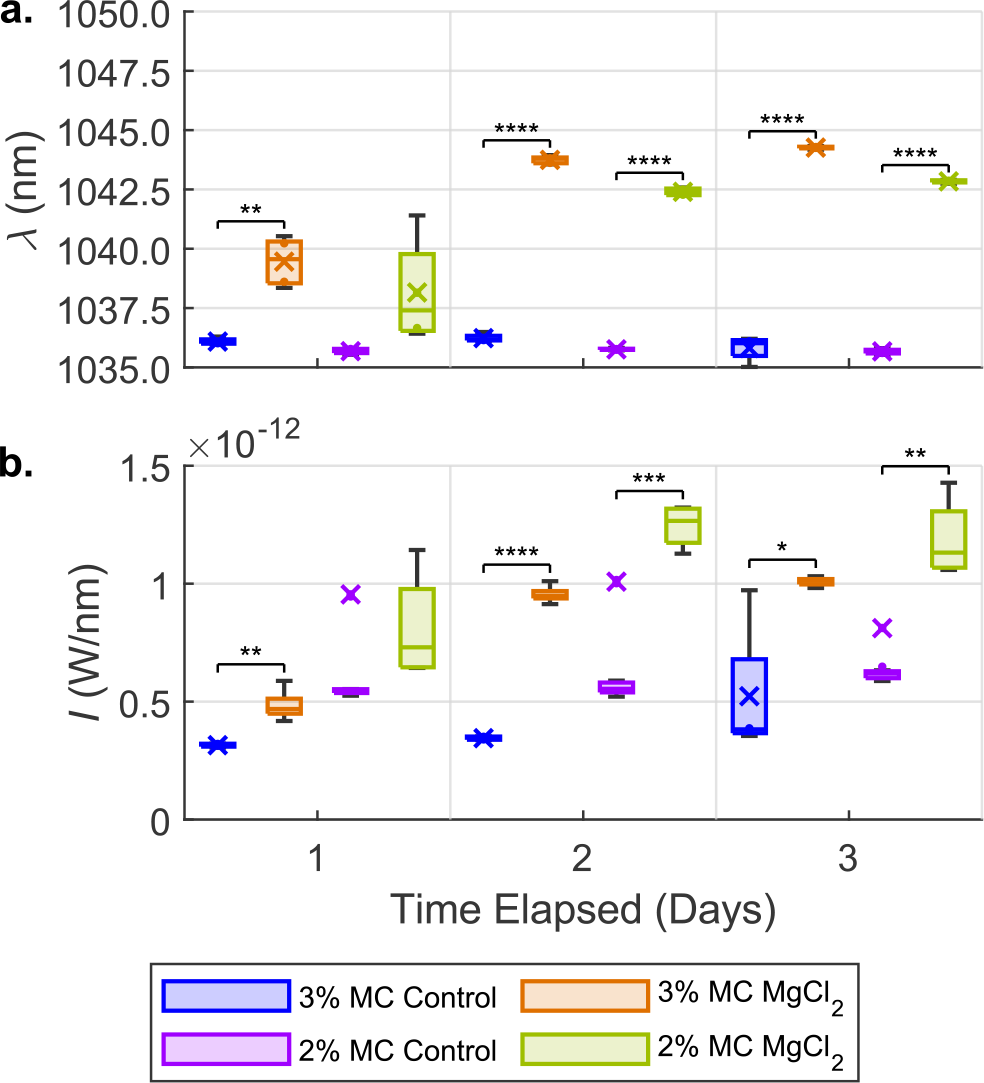
Sensor Response to MgCl_2_ in MC-based hydrogels. A) center wavelength and b) intensity. P-values for 3% MC shifts on days one through three were .001, 4 × 10^-10^, and 6 × 10^-5^ respectively; p-values for 2% MC on days two and three were 2 × 10^-6^ and 2 × 10^-12^ respectively. P-values for 3% MC intensity changes on days one through three were .004, 9 × 10^-7^, and .048 respectively; p-values for 2% MC on days two and three were 3 × 10^-4^ and .006 respectively. N = 4-5.

**Figure 5.**
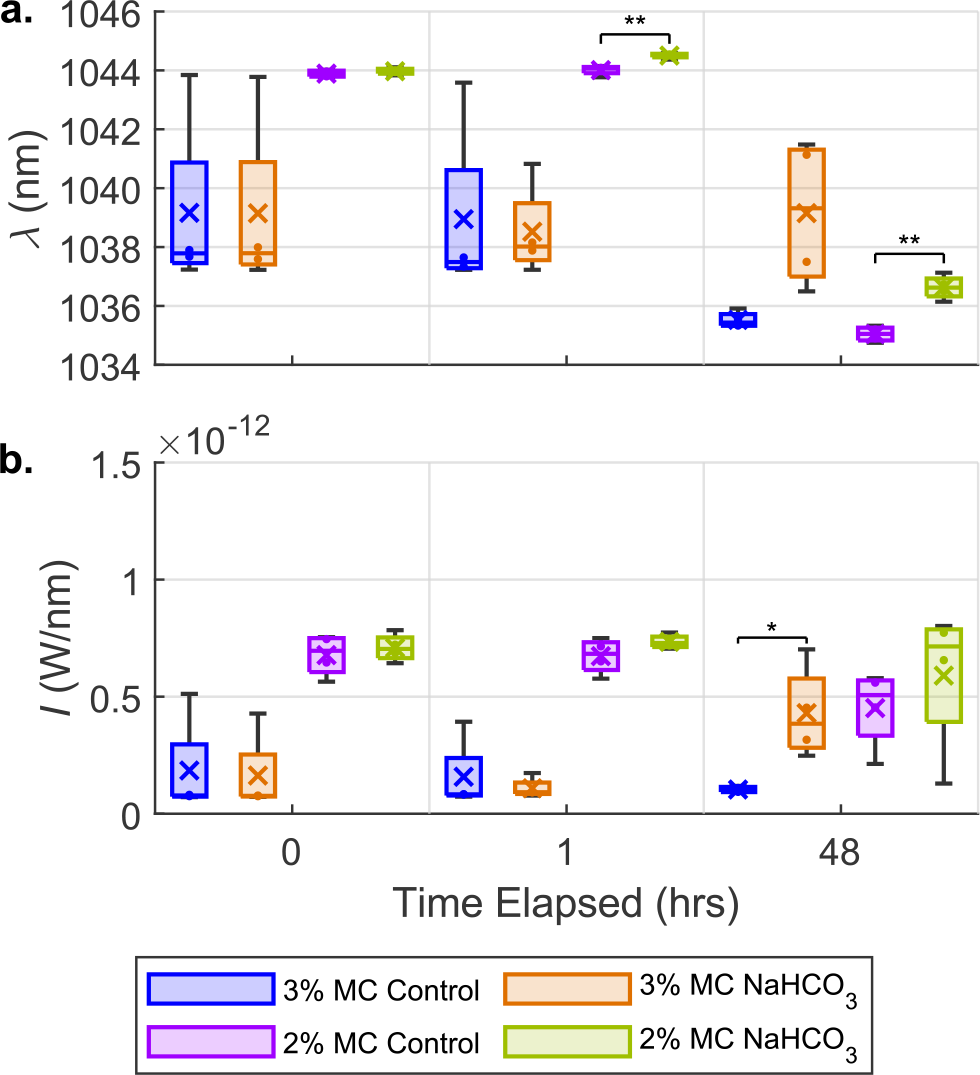
Sensor Response to NaHCO_3_ in MC-based hydrogels. A) center wavelength and b) intensity. The 2% sensor gel exhibited a slight but significant shift at hours one and 48 (p = .004 and, 001 respectively), whereas the 3% sensor gel system only showed a significant intensity-based response on the second day (p = .047). N = 3-4.

### MgCl_2_ Quantification in Methylcellulose Gels

Because the MC-SWCNT system demonstrated the most substantial responses and has the most desirable mechanical properties, we further explored this system to evaluate the breadth of MC’s potential as an in vivo sensor delivery platform. To evaluate the system’s capacity for analyte quantification across a range of concentrations, MC-SWCNT samples were topped with MgCl_2_ from 4 to 1000 µM. Concentrationdependent shift- and intensity-based responses were observed after two days (Figure 6) and fit to a Hill model. Interestingly, the intensity-based response was found to have higher sensitivity, with a fit-derived K_D_ of about 20 µM, compared to about 50 µM for the shift response. The decoupled nature of shifting and intensity responses may allow for analyte quantification across a broader range of concentrations than would otherwise be possible.^47^

**Figure 6.**
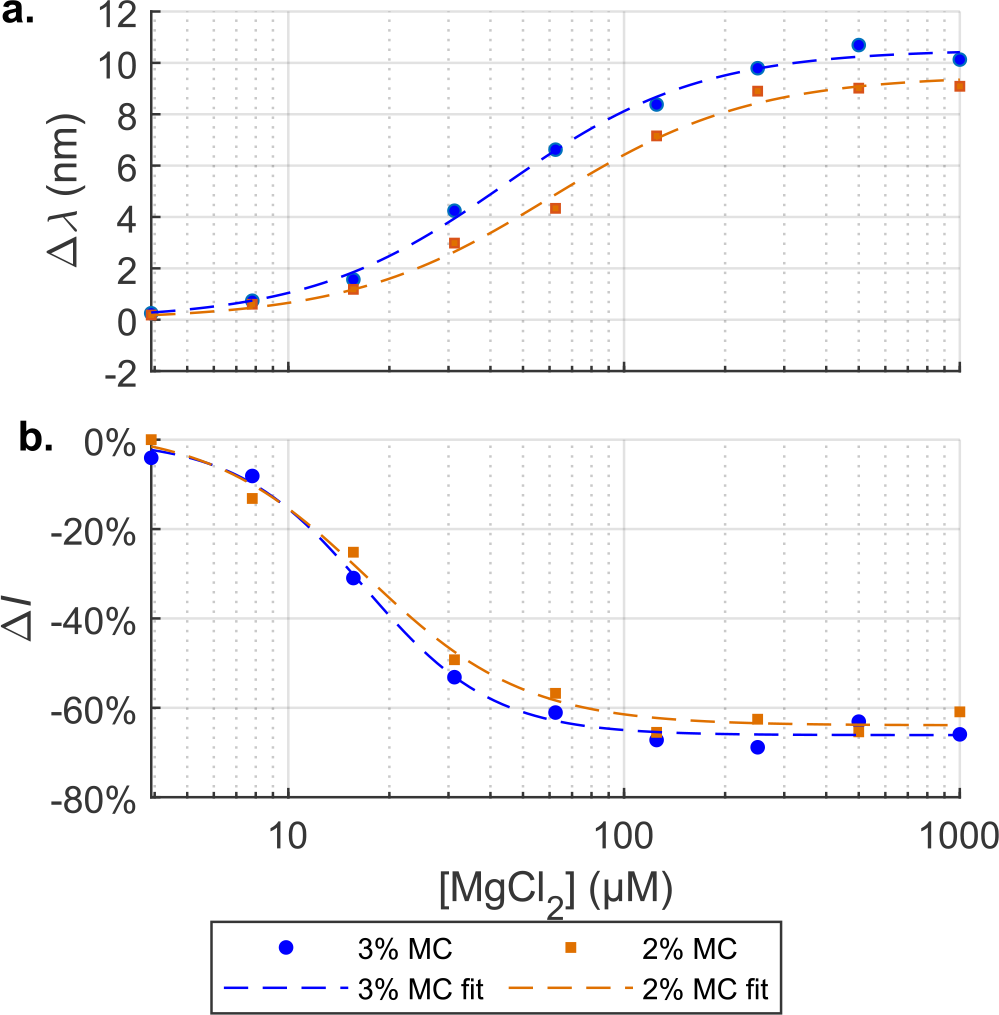
MgCl_2_ Quantification in MC Based Hydrogels. A) shift and b) intensity change. Fits use the Hill model; r^2^ ≥ 0.99 for all fits. Shift-derived K_D_ values are 44.1 and 60.3 µM for 3% and 2% gels respectively; intensity change-derived K_D_ values are 16.9 and 17.1 µM for 3% and 2% gels respectively.

### Detection of MgCl_2_ and Bovine Serum Albumin in Sulfonated Methylcellulose Gels

To potentially increase the response rate while maintaining the MC-SWCNT system’s mechanical properties, sulfonated versions were also investigated. As MC gels are largely hydrophobic, it was hypothesized that increasing polymer hydrophilicity by charge functionalization would better facilitate the diffusion of salts and proteins. Sulfonate groups were added to the MC polymers by replacing some of the methacrylate groups prior to gelation. This was accomplished by adapting a previously described modification of our gel preparation procedure.^48^ SWCNTs encapsulated in sulfonated 3% MC exhibited fluorescence across the entire 149-day duration of a stability assessment experiment (Figure S2). Following gelation of the 3% sulfonated methylcellulose formulation with encapsulated SWCNTs, we found that its viscoelastic properties were maintained in comparison to controls with minor, but nonsignificant reductions in both elasticity modulus and gelation time (Figure S3).

2% and 3% concentrations of the sulfonated methylcellulose (MC-SO_3_) gel with (GT)_15_-SWCNTs were evaluated for their response to MgCl_2_ (5 M) and bovine serum albumin (BSA, 600 µM). BSA was introduced as a representative large molecular weight globular protein, though it should be noted that noninvasive sensing of albumin has particularly important clinical implications in liver, renal, and cardiovascular disorders.^49^ Indeed, a SWNCT-based nanosensor paint has previously been reported for the pre-clinical detection of albuminuria.^50^ Both the sulfonated systems were found to have significant shift- and intensity-based responses to MgCl_2_ or BSA one day after incubation, an improved detection speed compared to the non-sulfonated devices (Figure 7). At this timepoint, the 3% gel-encapsulated SWCNTs responded to BSA with a shift and intensity change of +2.7 ± 0.6 and +660 ± 42% respectively; for the 2% system wavelength and intensity responses of +5.5 ± 0.7 and +360 ± 50% were observed. While wavelength shifts were maintained throughout the experiment, intensity differences moderated between days 2 and 6, largely due to increases in control intensity rather than diminution of analyte response. Following the success of these initial investigations into MC-SO_3_, an experiment was conducted to determine the effect of gel sulfonation on analyte detection. These experiments established that, as hypothesized, MC sulfonation improved BSA detection speed (Figure S4). Sulfonation was also shown to increase BSA response magnitude.

**Figure 7.**
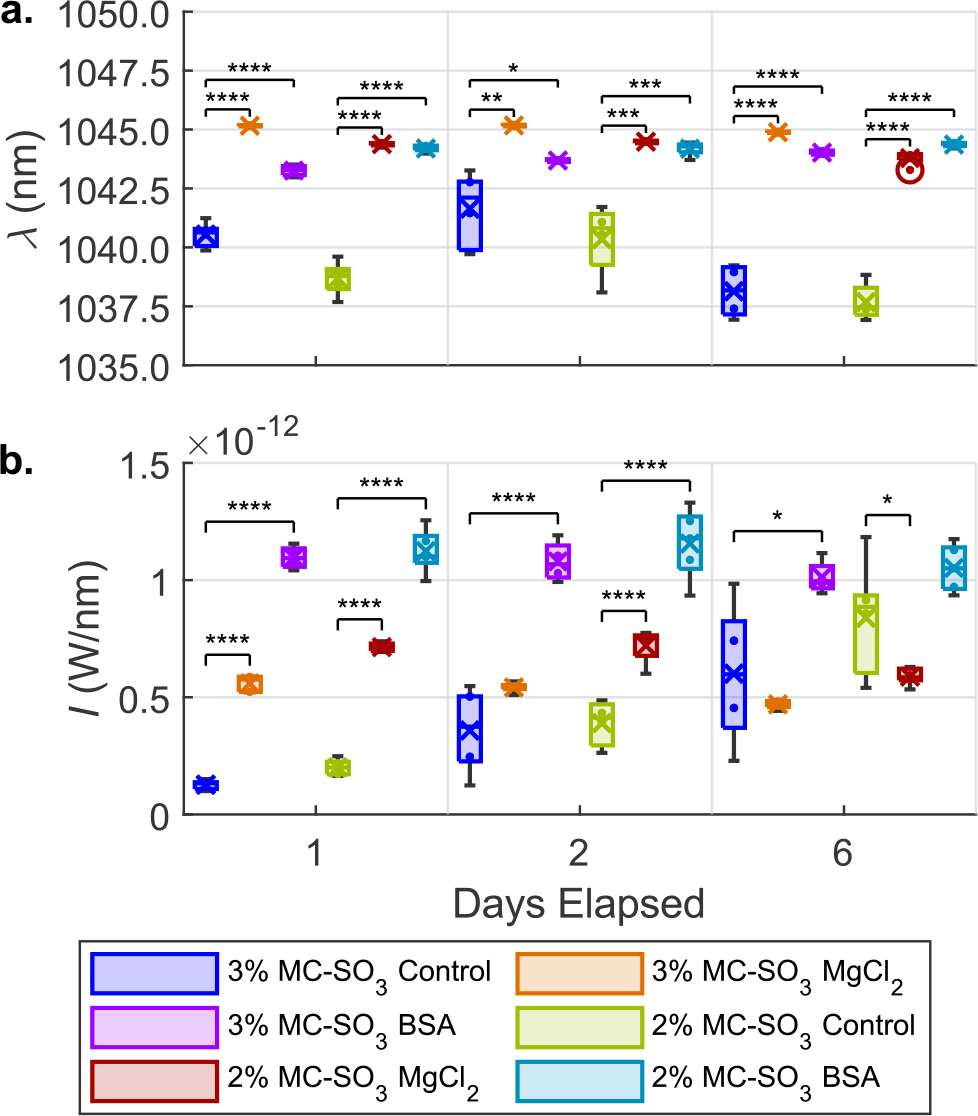
Sensor Response to MgCl_2_ and BSA in Sulfonated MC Hydrogels. A) center wavelength and b) intensity. Significant wavelength (p < .0001, Table S1) differences were observed for all groups on days 1 and 6; significant intensity differences (p < .0001, Table S2) were observed for all groups on days 1 and 2 (except for MgCl_2_ on day 2 in 2% MC-SO_3_). N ≥ 4.

BSA diffusion into MC gels was confirmed by a BCA assay (Figure S5). Analyte-containing topping solutions were removed and replaced with a solution of cellulase. Protein quantification was then performed on the degraded gels to determine the amount of BSA present, finding that gels exposed to BSA contained 130-250 µM of the protein, and that there were no clear differences in diffusion between the 2% and 3% gels.

We then evaluated the ability of the 3% MC-SO_3_-SWCNT system to maintain its BSA response over a two-week period. The 2% MC formulation was not used because sulfonation had a greater impact on response speed and the 3% concentration better approximates tissue. Significant wavelength- and intensity-based responses were observed across the entire experiment duration (Figure S6**Error! Reference source not found**.). We found that the wavelength-based response diminished across the experiment duration (from +8.9 ± 0.5 nm to +2.9 ± 1 nm), whereas the intensity response plateaued after three days (at +62% ± 10% of the control’s intensity). While the wavelength-based response can largely be attributed to red-shifting of the experimental group, the intensity-based response derives from intensity changes in the control group (Figure S7).

### Quantification of BSA and Doxorubicin in Sulfonated Methylcellulose Gels

Concentration-response curves were obtained in gel and solution to evaluate the effect of 3% MC-SO_3_ encapsulation on (GT)_15_-SWCNT response dynamics at 37 °C. Responses were measured at analyte concentrations from 10 nM to 10 mM. These experiments were executed using a high-throughput 96-well plate format, enabled by a custom-built near-IR plate-reader spectrophotometer. Responses were fit to a logistic trend (Equation S2). Here, we again investigated responses to albumin while separately introducing a new analyte, doxorubicin (DOX). DOX is a small molecule anthracycline chemotherapeutic used as a front-line therapy in several cancers.^21^ Our previous studies demonstrated SWCNT-based fluorescent sensors implanted within a dialysis membrane for DOX pharmacokinetic monitoring in mice.^21^

Whereas 3% MC-SO_3_ encapsulation increased BSA sensitivity (Figure 8) and response magnitude versus solution conditions, it also decreased DOX sensitivity (Figure 9). For BSA, the shift-derived K_D_ decreased by one to two orders of magnitude for every chirality except (8,7) (e.g., 920 to 16.7 µM for the (7,5) chirality, Table S4). Interestingly, intensity change-derived K_D_ values did not change appreciably (Table S5). In contrast, gel encapsulation increased the intensity change-derived DOX K_D_ by one to two orders of magnitude (e.g., 10.7 to 134. µM for the (7,5) chirality, Table S6). However, the MC-SO_3_ system better fit the expected trend across the tested concentration range, likely due to DOX aggregation at 10 µM in 1X PBS.^51^

**Figure 8.**
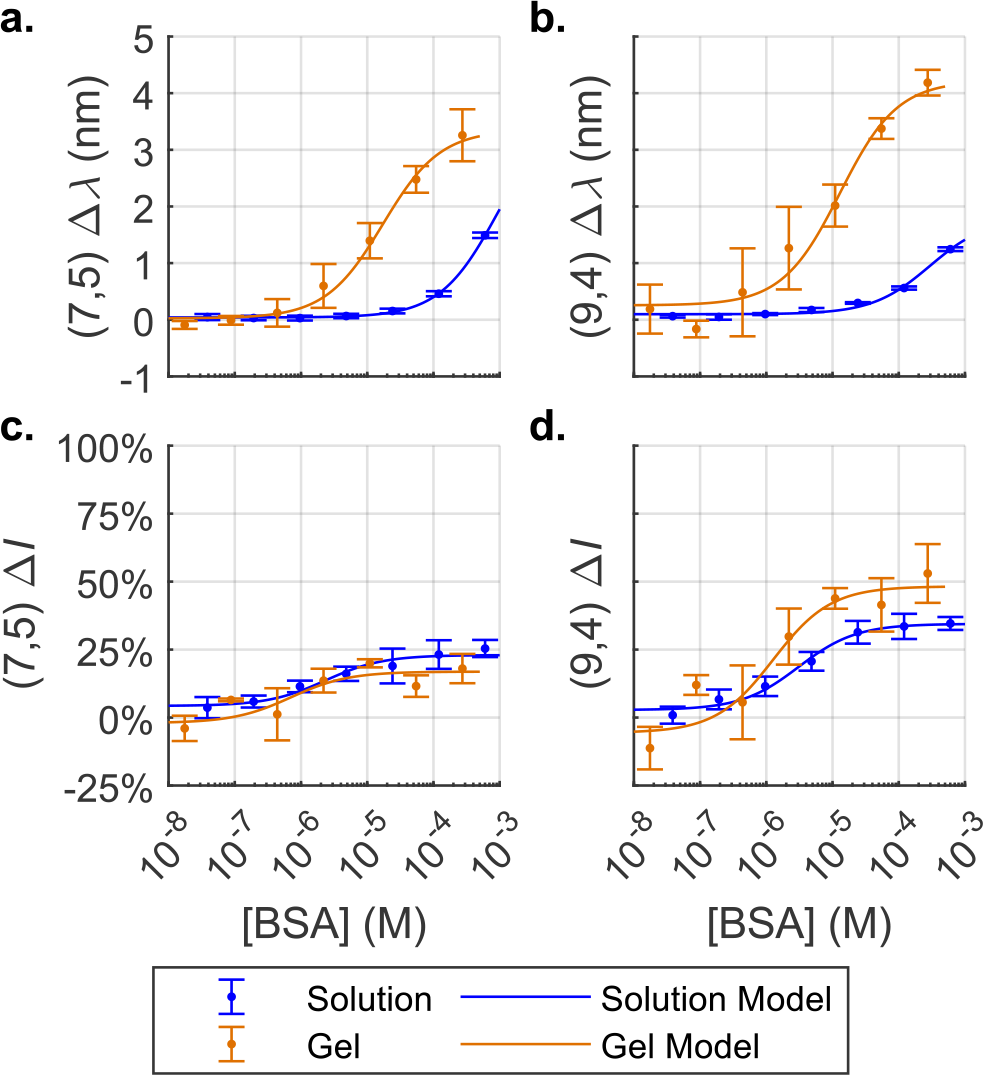
Quantification of BSA in 3% MC-SO_3_ Hydrogels and Solution. Shifts of a) (7,5) and b) (9,4) chiralities. Intensity changes of c) (7,5) and d) (9,4) chiralities. Data were fit to a logistic model (Equation S2); fit parameters/quality recorded in SI (Tables S4, S5).

**Figure 9.**
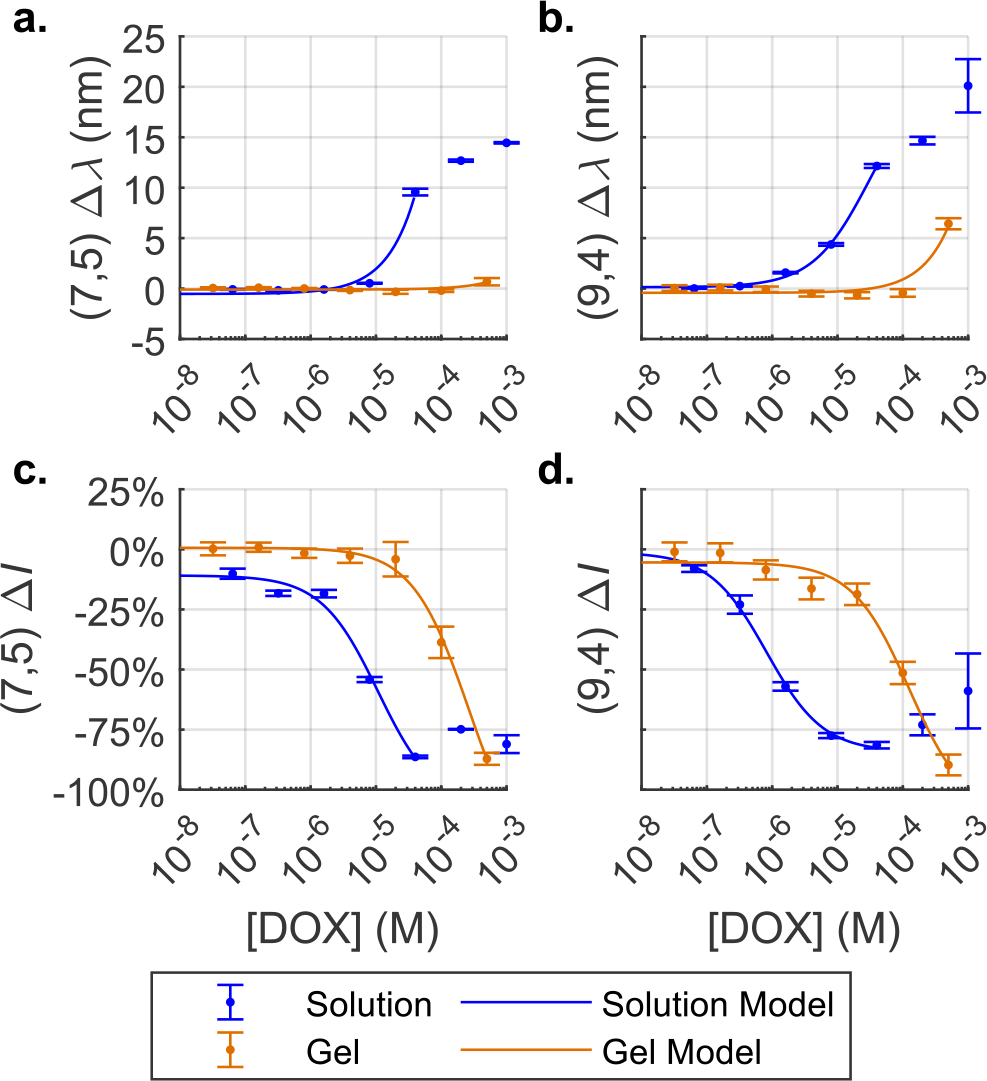
Quantification of Doxorubicin in 3% MC-SO_3_ Hydrogels and Solution. Shifts of a) (7,5) and b) (9,4) chiralities. Intensity changes of c) (7,5) and d) (9,4) chiralities. Data were fit to a logistic model (Equation S2); fit parameters/quality recorded in SI (Supplementary Table 6). For solution data, the model was not extended to concentrations which disobeyed the trend because of DOX aggregation.

For both analytes, response curve goodness-of-fit was decreased after gel encapsulation, likely because of variance in gel volumes expelled by the dual-barrel syringe (Tables S4-6). In the case of BSA, this increased variance is partially offset by an increased shift magnitude, leading to a higher signal-to-noise ratio. Response times also increased; whereas the maximum responses were achieved by the first hour in solution, the maximum BSA and DOX responses were observed after 24 and 48 hours respectively.

### Doxorubicin Detection in Vivo

Following in vitro validation of this system, we sought to detect the small molecule chemotherapeutic doxorubicin in live animals. First, we injected a mouse with MC-SO_3_-encapsulated SWCNTs to evaluate the stability of the device in vivo. Clear near-infrared fluorescence emanating from gel-encapsulated SWCNTs was observed on each day investigated, up to two months post-injection. Minimal variations in emission spectra were noted upon excitation at 655 and 730 nm (Figure 10).

**Figure 10.**
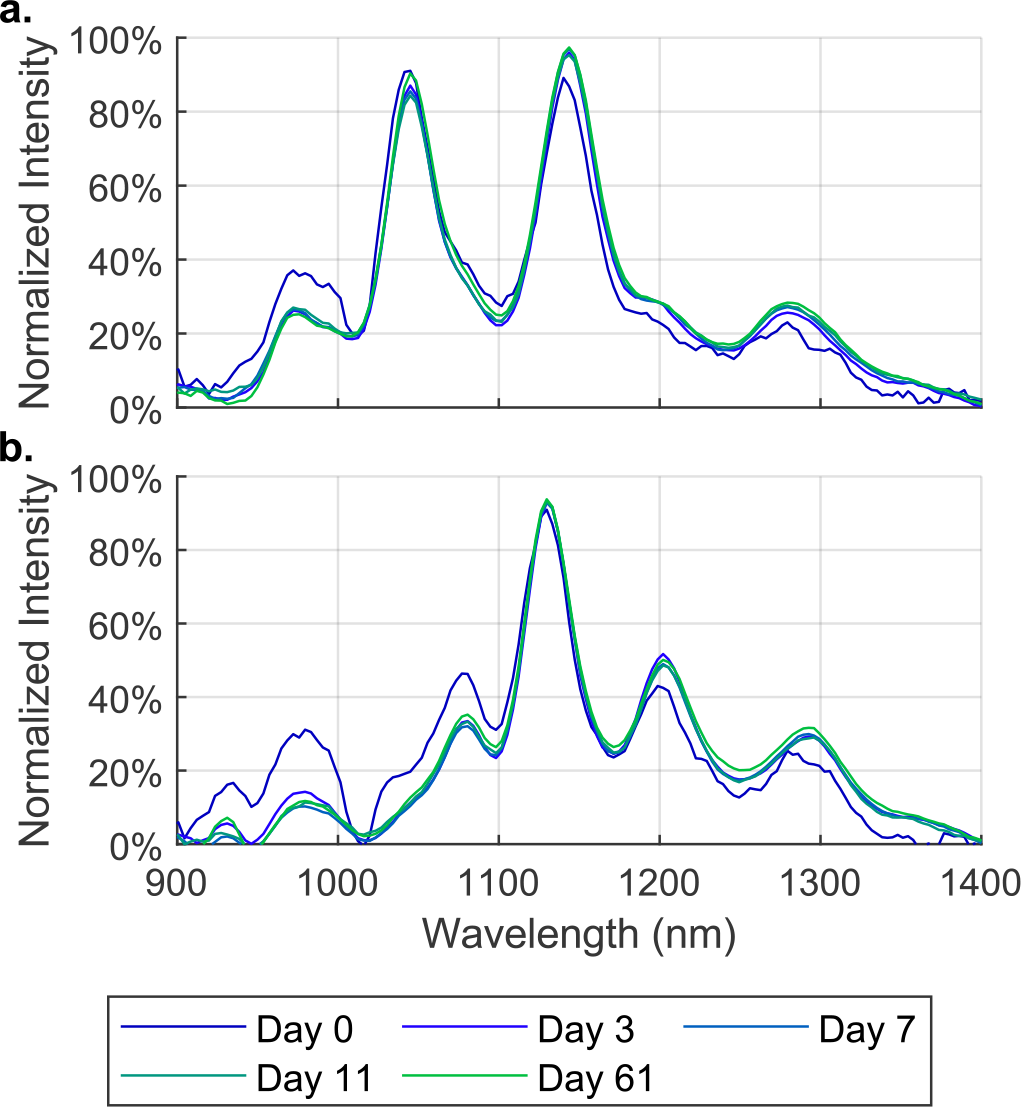
Fluorescence Spectra from (GT)_15_-SWCNT Encapsulated in 3% MC-SO_3_ Hydrogels Subcutaneously Injected Across 61 Days. a) 655 nm excitation b) 730 nm excitation.

To evaluate the function of the MC-SO_3_-SWCNT formulation in vivo, it was injected into six mice. After the implants were allowed to crosslink (approximately 15 minutes), the mice were anesthetized with isoflurane, and baseline spectra were acquired using an IRina NIR-II spectral probe with 655 and 730 nm excitation lasers coupled to an InGaAs detector. Mice were then subcutaneously dosed with DOX or 1X PBS as control at four sites around the implant location (n = 3). Fluorescence spectra were again acquired immediately, 10 minutes, 4 hours, 24 hours, and 48 hours post-dosing.

Analyses of these spectra showed that the implants exhibited a shifting-based response to DOX in vivo (**Figure 11**). DOX caused the (7,6) and (9,4) chiralities to shift (experimental minus control) by 1.3 ± 0.5 and 1.2 ± 0.6 nm respectively by the four-hour timepoint (**Figure 11**), whereas the relatively dim (8,6) chirality only showed a significant response after 48 hours (Figures S11, S12). Shifts at the 48-hour timepoint for the (7,6), (9,4), and (8,6) chiralities were, 1.7 ± 0.5, 0.6 ± 0.2 and 1.1 ± 0.3 nm respectively. The (7,5) chirality was also bright enough to be accurately fit, but exhibited no response (Figures S11, S12).

**Figure 11.**
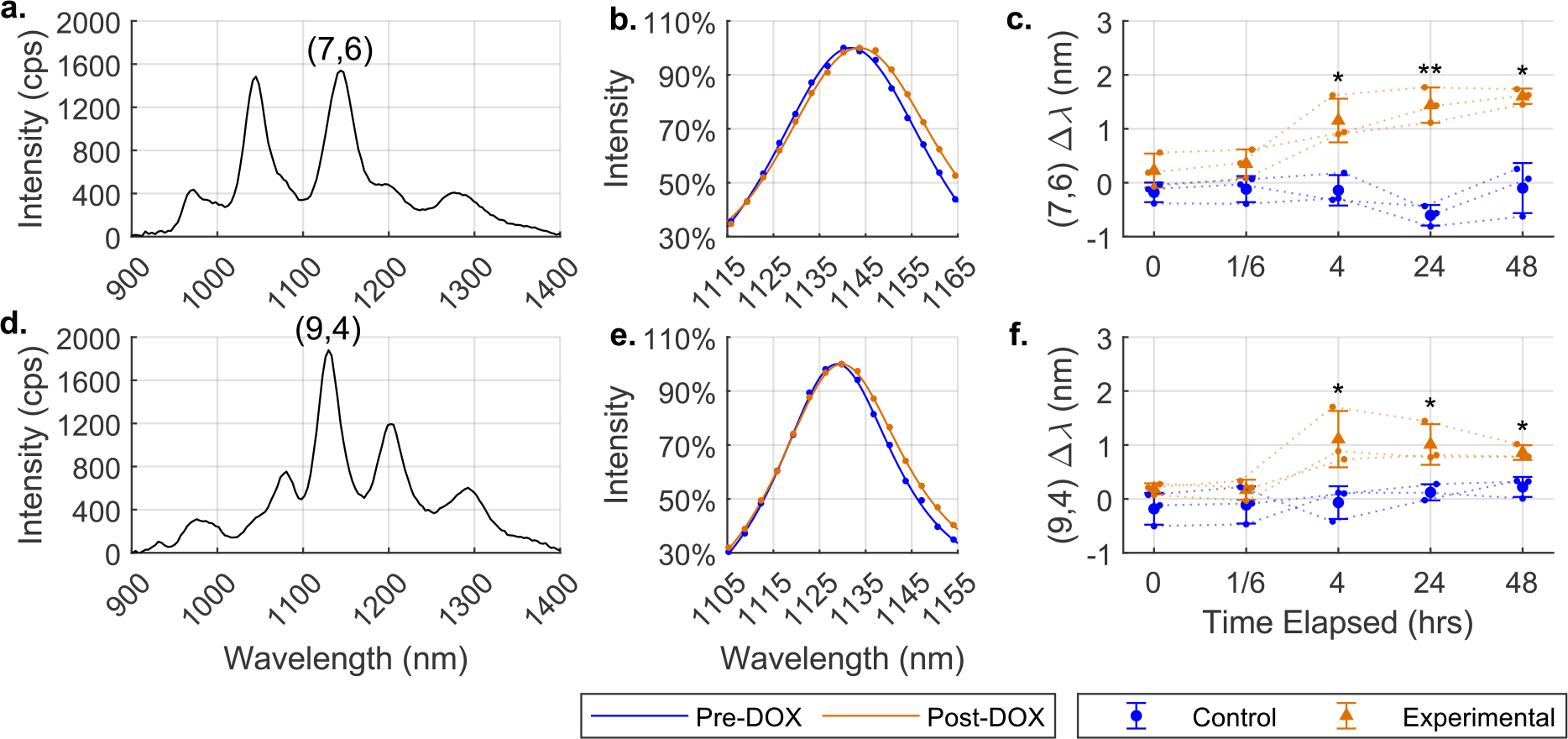
DOX Wavelength Response of Sensor Implants in Vivo. Representative autofluorescence-subtracted spectra of implants in vivo with a) 655 nm excitation and d) 730 nm excitation. Fluorescence peaks (data as dots, models as lines) from the same mouse just before and two days after DOX administration of b) (7,6) and e) (9,4) chiralities. Shifting over time of implants in control and experimental groups for c) (7,6) and d) (9,4) chiralities. N = 3. Error bars represent standard deviations. Significance of experimental shift versus control reported in Table S7.

While it was not possible to observe quenching because of high variation in both experimental and control group intensities (attributable to differences in sensor positioning across measurements), DOX did cause a ratiometric intensity response (Figure 12). In the experimental group, the (9,4) chirality got dimmer relative to the (7,6); the same effect was not observed in the control group.

While we have previously demonstrated in vivo DOX detection by (GT)_15_-SWCNTs implanted in dialysis membranes, we believe this novel approach utilizing an injectable hydrogel has better potential for clinical translation.^12^ It is likely that such an injectable, in situ stabilizing nanosensor formulation has substantially stronger potential due to its lower invasiveness.

**Figure 12.**
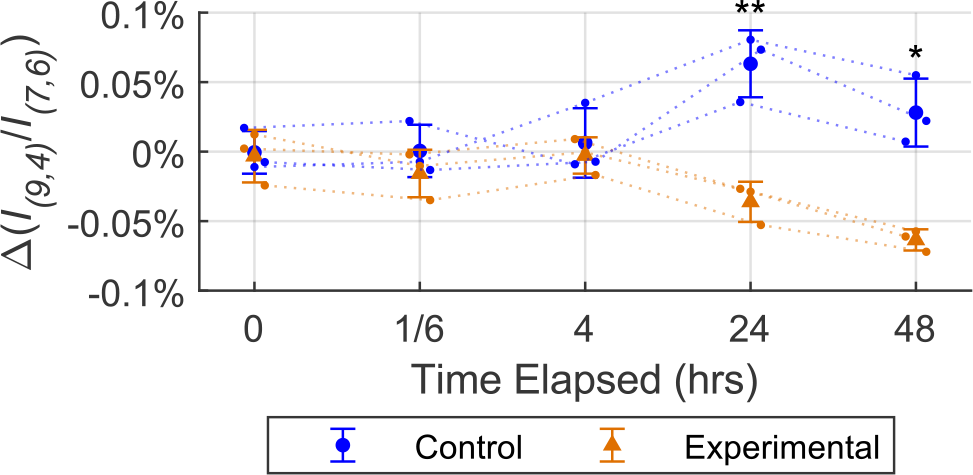
DOX Ratiometric Response of Sensor Implants in Vivo. Percent change in ratio of (9,4) fluorescence intensity to (7,6) fluorescence intensity. In the experimental group, the (9,4) chirality got dimmer relative to the (7,6) chirality. This relative dimming was not observed for the control group. p = .007 and .02 at hours 24 and 48 respectively.

## Conclusion

We found that 3% MC-SO_3_ is a functional platform for SWCNT-based biosensor encapsulation, demonstrating the potential of injectable hydrogels as a non-invasive means of implanting SWCNT biosensors. Beyond confirming that a salt (MgCl_2_) and a representative small-molecule drug (DOX) diffuse through the polymer matrix, we also observed that BSA, a 66.5 kDa protein, was able to permeate the gel and interact with SWCNTs. This confirms that the pore size of 3% MC-SO_3_ is not an obstacle to large biomarker diffusion. These experiments have also demonstrated that relatively precise analyte quantification is possible with this method of SWCNT encapsulation and delivery; logistic models of (GT)_15_-SWCNT fluorescence response as a function of analyte concentration showed R^2^ values as high as 0.99. For detection of the model globular protein analyte BSA, both response sensitivity and magnitude were increased in MC-SO_3_ compared to solution. The system also exhibits promising stability and functionality in vivo; implanted in mice, the sensors were shown to fluoresce for at least 61 days, and to discriminate mice injected with DOX from those injected with PBS as a control. We anticipate that further studies will continue to optimize this nanosensor delivery system for in vivo analyte detection and long-term compatibility for clinical use.

## Acknowledgments

The authors wish to acknowledge support of all members of the Williams and Nicoll Labs. This work was supported by NIH NIGMS R35GM142833 (RMW), a CCNY-MSKCC Partnership for Cancer Research Pilot award NIH NCI U54CA132378 and U54CA137788 (RMW and SBN), and a Stand Up to Cancer DISRUPT Pilot award (RMW and SBN).

## Notes

### Competing Interest Statement

The authors have declared no competing interest.

## References

(1) Nissler, R.; Ackermann, J.; Ma, C.; Kruss, S. Prospects of Fluorescent Single-Chirality Carbon Nanotube-Based Biosensors. Anal Chem 2022, 94 (28), 9941–9951. DOI: 10.1021/acs.analchem.2c01321 From NLM Medline.

(2) Weisman, R. B.; Bachilo, S. M. Dependence of Optical Transition Energies on Structure for Single-Walled Carbon Nanotubes in Aqueous Suspension: An Empirical Kataura Plot. Nano Letters 2003, 3 (9), 1235–1238. DOI: 10.1021/nl034428i (acccessed 2022-04-28T17:26:07).

(3) Hong, G.; Antaris, A. L.; Dai, H. Near-infrared fluorophores for biomedical imaging. Nature Biomedical Engineering 2017, 1 (1). DOI: 10.1038/s41551-016-0010.

(4) Boghossian, A. A.; Zhang, J.; Barone, P. W.; Reuel, N. F.; Kim, J. H.; Heller, D. A.; Ahn, J. H.; Hilmer, A. J.; Rwei, A.; Arkalgud, J. R.; et al. Near-infrared fluorescent sensors based on single-walled carbon nanotubes for life sciences applications. ChemSusChem 2011, 4 (7), 848–863. DOI: 10.1002/cssc.201100070 (acccessed 2021/04/15). From NLM Medline.

(5) Iverson, N. M.; Barone, P. W.; Shandell, M.; Trudel, L. J.; Sen, S.; Sen, F.; Ivanov, V.; Atolia, E.; Farias, E.; McNicholas, T. P.; et al. In vivo biosensing via tissue-localizable near-infrared-fluorescent single-walled carbon nanotubes. Nat Nanotechnol 2013, 8 (11), 873–880. DOI: 10.1038/nnano.2013.222 From NLM Medline.

(6) Koman, V. B.; Bakh, N. A.; Jin, X.; Nguyen, F. T.; Son, M.; Kozawa, D.; Lee, M. A.; Bisker, G.; Dong, J.; Strano, M. S. A wavelength-induced frequency filtering method for fluorescent nanosensors in vivo. Nat Nanotechnol 2022, 17 (6), 643–652. DOI: 10.1038/s41565-022-01136-x From NLM Medline.

(7) Gong, X.; Cho, S. Y.; Kuo, S.; Ogunlade, B.; Tso, K.; Salem, D. P.; Strano, M. S. Divalent Metal Cation Optical Sensing Using Single-Walled Carbon Nanotube Corona Phase Molecular Recognition. Anal Chem 2022, 94 (47), 16393–16401. DOI: 10.1021/acs.analchem.2c03648 From NLM Medline.

(8) Jin, H.; Jeng, E. S.; Heller, D. A.; Jena, P. V.; Kirmse, R.; Langowski, J.; Strano, M. S. Divalent ion and thermally induced DNA conformational polymorphism on single-walled carbon nanotubes. Macromolecules 2007, 40 (18), 6731–6739.

(9) Heller, D. A.; Jeng, E. S.; Yeung, T.-K.; Martinez, B. M.; Moll, A. E.; Gastala, J. B.; Strano, M. S. Optical detection of DNA conformational polymorphism on single-walled carbon nanotubes. Science 2006, 311 (5760), 508–511.

(10) Zhang, J.; Boghossian, A. A.; Barone, P. W.; Rwei, A.; Kim, J. H.; Lin, D.; Heller, D. A.; Hilmer, A. J.; Nair, N.; Reuel, N. F.; et al. Single molecule detection of nitric oxide enabled by d(AT)15 DNA adsorbed to near infrared fluorescent single-walled carbon nanotubes. J Am Chem Soc 2011, 133 (3), 567–581. DOI: 10.1021/ja1084942 From NLM Medline.

(11) Zubkovs, V.; Wang, H.; Schuergers, N.; Weninger, A.; Glieder, A.; Cattaneo, S.; Boghossian, A. A. Bioengineering a glucose oxidase nanosensor for near-infrared continuous glucose monitoring. Nanoscale Adv 2022, 4 (11), 2420–2427. DOI: 10.1039/d2na00092j From NLM PubMed-not-MEDLINE.

(12) Harvey, J. D.; Williams, R. M.; Tully, K. M.; Baker, H. A.; Shamay, Y.; Heller, D. A. An in Vivo Nanosensor Measures Compartmental Doxorubicin Exposure. Nano Lett 2019, 19 (7), 4343–4354. DOI: 10.1021/acs.nanolett.9b00956 From NLM Medline.

(13) Son, M.; Mehra, P.; Nguyen, F. T.; Jin, X.; Koman, V. B.; Gong, X.; Lee, M. A.; Bakh, N. A.; Strano, M. S. Molecular Recognition and In Vivo Detection of Temozolomide and 5-Aminoimidazole-4-carboxamide for Glioblastoma Using Near-Infrared Fluorescent Carbon Nanotube Sensors. ACS Nano 2023, 17 (1), 240–250. DOI: 10.1021/acsnano.2c07264 From NLM Medline.

(14) Ang, M. C.; Dhar, N.; Khong, D. T.; Lew, T. T. S.; Park, M.; Sarangapani, S.; Cui, J.; Dehadrai, A.; Singh, G. P.; Chan-Park, M. B.; et al. Nanosensor Detection of Synthetic Auxins In Planta using Corona Phase Molecular Recognition. ACS Sens 2021, 6 (8), 3032–3046. DOI: 10.1021/acssensors.1c01022 From NLM Medline.

(15) Bisker, G.; Bakh, N. A.; Lee, M. A.; Ahn, J.; Park, M.; O’Connell, E. B.; Iverson, N. M.; Strano, M. S. Insulin Detection Using a Corona Phase Molecular Recognition Site on Single-Walled Carbon Nanotubes. ACS Sens 2018, 3 (2), 367–377. DOI: 10.1021/acssensors.7b00788 From NLM Medline.

(16) Lee, M. A.; Wang, S.; Jin, X.; Bakh, N. A.; Nguyen, F. T.; Dong, J.; Silmore, K. S.; Gong, X.; Pham, C.; Jones, K. K.; et al. Implantable Nanosensors for Human Steroid Hormone Sensing In Vivo Using a Self-Templating Corona Phase Molecular Recognition. Adv Healthc Mater 2020, 9 (21), e2000429. DOI: 10.1002/adhm.202000429 From NLM Medline.

(17) Zhang, J.; Landry, M. P.; Barone, P. W.; Kim, J. H.; Lin, S.; Ulissi, Z. W.; Lin, D.; Mu, B.; Boghossian, A. A.; Hilmer, A. J.; et al. Molecular recognition using corona phase complexes made of synthetic polymers adsorbed on carbon nanotubes. Nat Nanotechnol 2013, 8 (12), 959–968. DOI: 10.1038/nnano.2013.236 From NLM Medline.

(18) Kruss, S.; Landry, M. P.; Vander Ende, E.; Lima, B. M.; Reuel, N. F.; Zhang, J.; Nelson, J.; Mu, B.; Hilmer, A.; Strano, M. Neurotransmitter detection using corona phase molecular recognition on fluorescent single-walled carbon nanotube sensors. J Am Chem Soc 2014, 136 (2), 713–724. DOI: 10.1021/ja410433b From NLM Medline.

(19) Dinarvand, M.; Neubert, E.; Meyer, D.; Selvaggio, G.; Mann, F. A.; Erpenbeck, L.; Kruss, S. Near-Infrared Imaging of Serotonin Release from Cells with Fluorescent Nanosensors. Nano Lett 2019, 19 (9), 6604–6611. DOI: 10.1021/acs.nanolett.9b02865 From NLM Medline.

(20) Kelich, P.; Jeong, S.; Navarro, N.; Adams, J.; Sun, X.; Zhao, H.; Landry, M. P.; Vukovic, L. Discovery of DNA-Carbon Nanotube Sensors for Serotonin with Machine Learning and Near-infrared Fluorescence Spectroscopy. ACS Nano 2022, 16 (1), 736–745. DOI: 10.1021/acsnano.1c08271 From NLM Medline.

(21) Harvey, J. D.; Jena, P. V.; Baker, H. A.; Zerze, G. H.; Williams, R. M.; Galassi, T. V.; Roxbury, D.; Mittal, J.; Heller, D. A. A Carbon Nanotube Reporter of miRNA Hybridization Events In Vivo. Nat Biomed Eng 2017, 1 (4), 0041. DOI: 10.1038/s41551-017-0041 From NLM PubMed-not-MEDLINE.

(22) Bisker, G.; Dong, J.; Park, H. D.; Iverson, N. M.; Ahn, J.; Nelson, J. T.; Landry, M. P.; Kruss, S.; Strano, M. S. Protein-targeted corona phase molecular recognition. Nat Commun 2016, 7 (1), 10241. DOI: 10.1038/ncomms10241 From NLM Medline.

(23) Lee, K.; Nojoomi, A.; Jeon, J.; Lee, C. Y.; Yum, K. Near-Infrared Fluorescence Modulation of Refolded DNA Aptamer-Functionalized Single-Walled Carbon Nanotubes for Optical Sensing. ACS Applied Nano Materials 2018, 1 (9), 5327–5336. DOI: 10.1021/acsanm.8b01377.

(24) Antman-Passig, M.; Wong, E.; Frost, G. R.; Cupo, C.; Shah, J.; Agustinus, A.; Chen, Z.; Mancinelli, C.; Kamel, M.; Li, T.; et al. Optical Nanosensor for Intracellular and Intracranial Detection of Amyloid-Beta. ACS Nano 2022, 16 (5), 7269–7283. DOI: 10.1021/acsnano.2c00054 From NLM Medline.

(25) Jin, X.; Lee, M. A.; Gong, X.; Koman, V. B.; Lundberg, D. J.; Wang, S.; Bakh, N. A.; Park, M.; Dong, J. I.; Kozawa, D.; et al. Corona Phase Molecular Recognition of the Interleukin-6 (IL-6) Family of Cytokines Using nIR Fluorescent Single-Walled Carbon Nanotubes. ACS Applied Nano Materials 2023, 6 (11), 9791–9804. DOI: 10.1021/acsanm.3c01525.

(26) Jena, P. V.; Roxbury, D.; Galassi, T. V.; Akkari, L.; Horoszko, C. P.; Iaea, D. B.; Budhathoki-Uprety, J.; Pipalia, N.; Haka, A. S.; Harvey, J. D.; et al. A Carbon Nanotube Optical Reporter Maps Endolysosomal Lipid Flux. ACS Nano 2017, 11 (11), 10689–10703. DOI: 10.1021/acsnano.7b04743 From NLM Medline.

(27) Zhang, J.; Kruss, S.; Hilmer, A. J.; Shimizu, S.; Schmois, Z.; De La Cruz, F.; Barone, P. W.; Reuel, N. F.; Heller, D. A.; Strano, M. S. A rapid, direct, quantitative, and label-free detector of cardiac biomarker troponin T using near-infrared fluorescent single-walled carbon nanotube sensors. Adv Healthc Mater 2014, 3 (3), 412–423. DOI: 10.1002/adhm.201300033 From NLM Medline.

(28) Williams, R. M.; Lee, C.; Galassi, T. V.; Harvey, J. D.; Leicher, R.; Sirenko, M.; Dorso, M. A.; Shah, J.; Olvera, N.; Dao, F.; et al. Noninvasive ovarian cancer biomarker detection via an optical nanosensor implant. Sci Adv 2018, 4 (4), eaaq1090. DOI: 10.1126/sciadv.aaq1090 From NLM Medline.

(29) Williams, R. M.; Lee, C.; Heller, D. A. A Fluorescent Carbon Nanotube Sensor Detects the Metastatic Prostate Cancer Biomarker uPA. ACS Sens 2018, 3 (9), 1838–1845. DOI: 10.1021/acssensors.8b00631 From NLM Medline.

(30) Ackermann, J.; Metternich, J. T.; Herbertz, S.; Kruss, S. Biosensing with Fluorescent Carbon Nanotubes. Angew Chem Int Ed Engl 2022, 61 (18), e202112372. DOI: 10.1002/anie.202112372 From NLM Medline.

(31) Galassi, T. V.; Jena, P. V.; Shah, J.; Ao, G.; Molitor, E.; Bram, Y.; Frankel, A.; Park, J.; Jessurun, J.; Ory, D. S.; et al. An optical nanoreporter of endolysosomal lipid accumulation reveals enduring effects of diet on hepatic macrophages in vivo. Sci Transl Med 2018, 10 (461). DOI: 10.1126/scitranslmed.aar2680 From NLM Medline.

(32) Galassi, T. V.; Antman-Passig, M.; Yaari, Z.; Jessurun, J.; Schwartz, R. E.; Heller, D. A. Long-term in vivo biocompatibility of single-walled carbon nanotubes. PLoS One 2020, 15 (5), e0226791. DOI: 10.1371/journal.pone.0226791 (acccessed 2022-09-19T20:10:12). From NLM Medline.

(33) Hofferber, E.; Meier, J.; Herrera, N.; Stapleton, J.; Calkins, C.; Iverson, N. Detection of single walled carbon nanotube based sensors in a large mammal. Nanomedicine 2022, 40, 102489. DOI: 10.1016/j.nano.2021.102489 From NLM Medline.

(34) Bakh, N. A.; Gong, X.; Lee, M. A.; Jin, X.; Koman, V. B.; Park, M.; Nguyen, F. T.; Strano, M. S. Transcutaneous Measurement of Essential Vitamins Using Near-Infrared Fluorescent Single-Walled Carbon Nanotube Sensors. Small 2021, 17 (31), e2100540. DOI: 10.1002/smll.202100540 From NLM Medline.

(35) Lee, M. A.; Jin, X.; Muthupalani, S.; Bakh, N. A.; Gong, X.; Strano, M. S. In-Vivo fluorescent nanosensor implants based on hydrogel-encapsulation: investigating the inflammation and the foreign-body response. J Nanobiotechnology 2023, 21 (1), 133. DOI: 10.1186/s12951-023-01873-8 From NLM Medline.

(36) Lee, M. A.; Nguyen, F. T.; Scott, K.; Chan, N. Y. L.; Bakh, N. A.; Jones, K. K.; Pham, C.; Garcia-Salinas, P.; Garcia-Parraga, D.; Fahlman, A.; et al. Implanted Nanosensors in Marine Organisms for Physiological Biologging: Design, Feasibility, and Species Variability. ACS Sens 2019, 4 (1), 32–43. DOI: 10.1021/acssensors.8b00538 From NLM Medline.

(37) Correa, S.; Grosskopf, A. K.; Lopez Hernandez, H.; Chan, D.; Yu, A. C.; Stapleton, L. M.; Appel, E. A. Translational Applications of Hydrogels. Chem Rev 2021, 121 (18), 11385–11457. DOI: 10.1021/acs.chemrev.0c01177 From NLM Medline.

(38) Hofferber, E.; Meier, J.; Herrera, N.; Stapleton, J.; Ney, K.; Francis, B.; Calkins, C.; Iverson, N. Novel methods to extract and quantify sensors based on single wall carbon nanotube fluorescence from animal tissue and hydrogel-based platforms. Methods Appl Fluoresc 2021, 9 (2), 025005. DOI: 10.1088/2050-6120/abea07 From NLM Medline.

(39) Gold, G. T.; Varma, D. M.; Taub, P. J.; Nicoll, S. B. Development of crosslinked methylcellulose hydrogels for soft tissue augmentation using an ammonium persulfate-ascorbic acid redox system. Carbohydr Polym 2015, 134, 497–507. DOI: 10.1016/j.carbpol.2015.07.101 From NLM Medline.

(40) Chalanqui, M. J.; Pentlavalli, S.; McCrudden, C.; Chambers, P.; Ziminska, M.; Dunne, N.; McCarthy, H. O. Influence of alginate backbone on efficacy of thermo-responsive alginate-g-P(NIPAAm) hydrogel as a vehicle for sustained and controlled gene delivery. Mater Sci Eng C Mater Biol Appl 2019, 95, 409–421. DOI: 10.1016/j.msec.2017.09.003 From NLM Medline.

(41) Qin, H.; Wang, J.; Wang, T.; Gao, X.; Wan, Q.; Pei, X. Preparation and Characterization of Chitosan/beta-Glycerophosphate Thermal-Sensitive Hydrogel Reinforced by Graphene Oxide. Front Chem 2018, 6, 565. DOI: 10.3389/fchem.2018.00565 From NLM PubMed-not-MEDLINE.

(42) Cohen, Z.; Parveen, S.; Williams, R. M. Optimization of ssDNA-SWCNT Ultracentrifugation via Efficacy Measurements. ECS Journal of Solid State Science and Technology 2022, 11 (10). DOI: 10.1149/2162-8777/ac9929.

(43) Bachilo, S. M.; Strano, M. S.; Kittrell, C.; Hauge, R. H.; Smalley, R. E.; Weisman, R. B. Structure-assigned optical spectra of single-walled carbon nanotubes. Science 2002, 298 (5602), 2361–2366. DOI: 10.1126/science.1078727 From NLM PubMed-not-MEDLINE.

(44) Larsen, B. A.; Deria, P.; Holt, J. M.; Stanton, I. N.; Heben, M. J.; Therien, M. J.; Blackburn, J. L. Effect of solvent polarity and electrophilicity on quantum yields and solvatochromic shifts of single-walled carbon nanotube photoluminescence. J Am Chem Soc 2012, 134 (30), 12485–12491. DOI: 10.1021/ja2114618 From NLM PubMed-not-MEDLINE.

(45) Choi, J. H.; Strano, M. S. Solvatochromism in single-walled carbon nanotubes. Applied Physics Letters 2007, 90 (22). DOI: 10.1063/1.2745228.

(46) Varghese, S. A.; Rangappa, S. M.; Siengchin, S.; Parameswaranpillai, J. Natural polymers and the hydrogels prepared from them. In Hydrogels Based on Natural Polymers, 2020; pp 17–47.

(47) Card, M.; Alejandro, R.; Roxbury, D. Decoupling Individual Optical Nanosensor Responses Using a Spin-Coated Hydrogel Platform. ACS Appl Mater Interfaces 2023, 15 (1), 1772–1783. DOI: 10.1021/acsami.2c16596 From NLM Medline.

(48) Lin, H. A. Development and Functional Evaluation of Injectable Cellulose-based Hydrogels as Nucleus Pulposus Replacements for Intervertebral Disc Repair. Ph.D., The City College of New York, United States -- New York, 2018. https://www.proquest.com/docview/2091422462.

(49) Tabata, F.; Wada, Y.; Kawakami, S.; Miyaji, K. Serum Albumin Redox States: More Than Oxidative Stress Biomarker. Antioxidants (Basel) 2021, 10 (4). DOI: 10.3390/antiox10040503 From NLM PubMed-not-MEDLINE.

(50) Budhathoki-Uprety, J.; Shah, J.; Korsen, J. A.; Wayne, A. E.; Galassi, T. V.; Cohen, J. R.; Harvey, J. D.; Jena, P. V.; Ramanathan, L. V.; Jaimes, E. A.; et al. Synthetic molecular recognition nanosensor paint for microalbuminuria. Nat Commun 2019, 10 (1), 3605. DOI: 10.1038/s41467-019-11583-1 From NLM Medline.

(51) Zhu, L.; Yang, S.; Qu, X.; Zhu, F.; Liang, Y.; Liang, F.; Wang, Q.; Li, J.; Li, Z.; Yang, Z. Fibril-shaped aggregates of doxorubicin with poly-l-lysine and its derivative. Polym. Chem. 2014, 5 (19), 5700–5706. DOI: 10.1039/c4py00686k.

