## Supplementary Information for "Noninvasive Injectable Optical Nanosensor-Hydrogel Hybrids Detect Doxorubicin in Living Mice"

### Supplementary Methods

#### Rheological Characterization of Sulfonated Methylcellulose Gels

Rheological measurements were performed as previously described.[1] Characterization was executed at room temperature (n = 3-5) with an AR2000ex Rheometer (TA Instruments, New Castle, DE) using cone and plate geometry (2°, 40 mm). Parameters (0.5% strain, 1 Hz) were selected based on previous work. The gelation time was defined as the delay between initiator mixing and the timepoint when the derivative of *G'* (the elasticity modulus) was less than 1.

#### Stability of Sulfonated Methylcellulose Gel-Encapsulated SWCNT Over 149 days

Sterile 3% MC-$SO_3$ gels containing $(GT)_{15}$-SWCNTs (1 mg/L) were cast into cuvettes (250-500 mg gel per cuvette, 5 cuvettes total). After 30 minutes, cuvettes were topped with 1X PBS (1 mL). Spectra were acquired after 1, 7, 37, 45, 51, 58, 135, and 149 days. days. Between measurements, cuvettes were stored in an incubator at 37 °C.

#### BCA Assay

2% and 3% MC gels containing $(GT)_{15}$-SWCNTs (1 mg/L) were cast into pre-weighed cuvettes (250-500 mg gel per cuvette). After 30 minutes, cuvettes were topped with 1X PBS or BSA (600 µM) in 1X PBS (n = 5 – 6). Topping solution volumes were 1 µL per 1 mg gel. Cuvettes were stored in an incubator at 37 °C for two days before topping solutions were removed and replaced with cellulase (1000 U/L, 0.4739 µL per mg gel) in a pH 5 citrate buffer as previously described.[1] Cuvettes were again incubated at 37 °C overnight. A Micro BCA Protein Assay (Thermo Fisher Scientific, Waltham, MA, USA) was then used to quantify the BSA content of the degraded gels according to manufacturer instructions.

| [MC] | 2% | 3% |
|---|---|---|
| Control | 6 | 5 |
| BSA | 5 | 6 |

#### Effect of Sulfonation on BSA and DOX Responses

3% MC and 3% MC-$SO_3$ gels containing $(GT)_{15}$-SWCNTs (1 mg/L) were cast into a 96-well plate (150 ± 50 µL per well, 24 wells). After 30 minutes crosslinking time, fluorescence spectra were acquired using a ClaIR IR plate-reader. Gels were then topped with 100 µL 1X PBS, BSA (300 µM) in 1X PBS, or DOX (100 µM) in 1X PBS (4 wells per concentration/sulfonation combination). More fluorescence spectra were acquired after 10 minutes, 3 hours, 6 hours, 24 hours, 48 hours, and 72 hours. The plate was stored at 37 °C in an incubator between measurements.

### Supplementary Figures

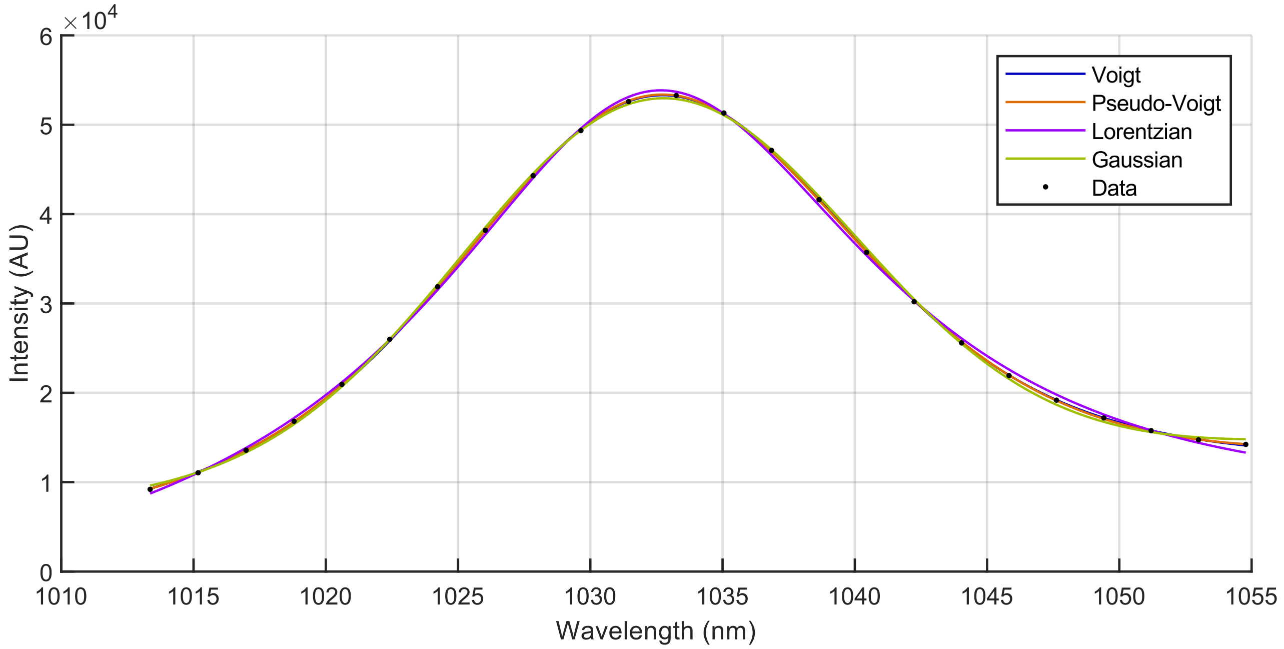


**Figure S1. Peak-Fitting Model Comparison.** Voigt, pseudo-Voigt, Lorentzian, and Gaussian fits of (7,5) $E_{11}$ peak. All models produce fits with sufficiently high $R^2$ values, but Voigt and pseudo-Voigt fits have greater agreement with data near center tails. Adjusted $R^2$ values for Voigt, Pseudo-Voigt, Lorentzian, and Gaussian models are 0.99996, 0.99996, 0.99892, and 0.99943 respectively.

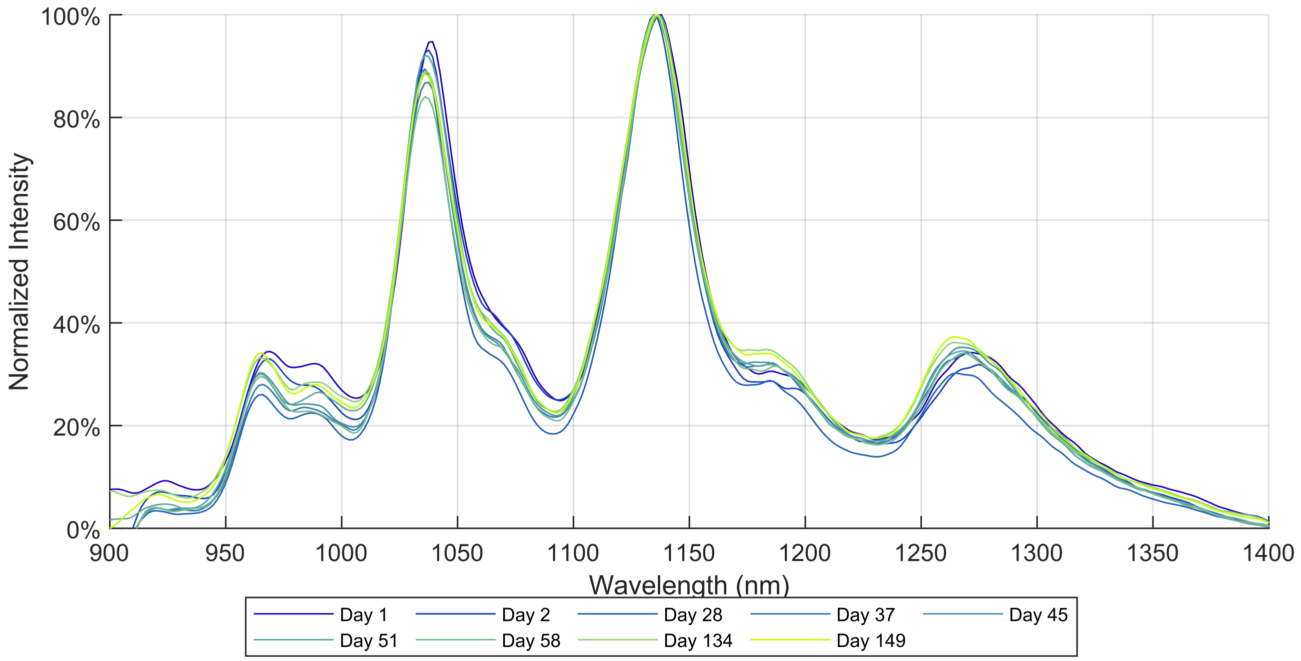

**Figure S2 Fluorescence Spectra of (GT)$_{15}$-SWCNTs in 3% MC-SO$_3$ Across 149 Days.**

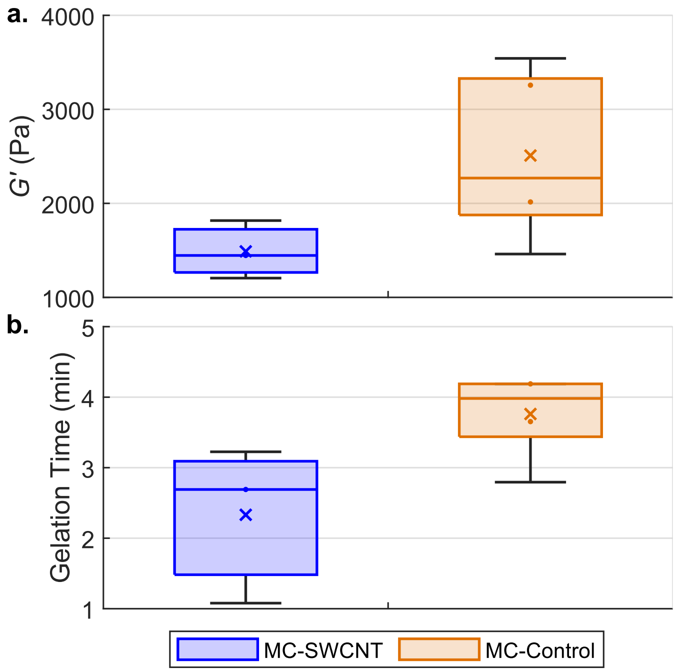


**Figure S3. Rheological comparison of 3% MC-SO$_3$ Gels with and Without** Sensors**.** a) elasticity modulus and b) gelation time, defined as point where the derivative of the elasticity modulus < 1 Pa. No significant differences were observed.

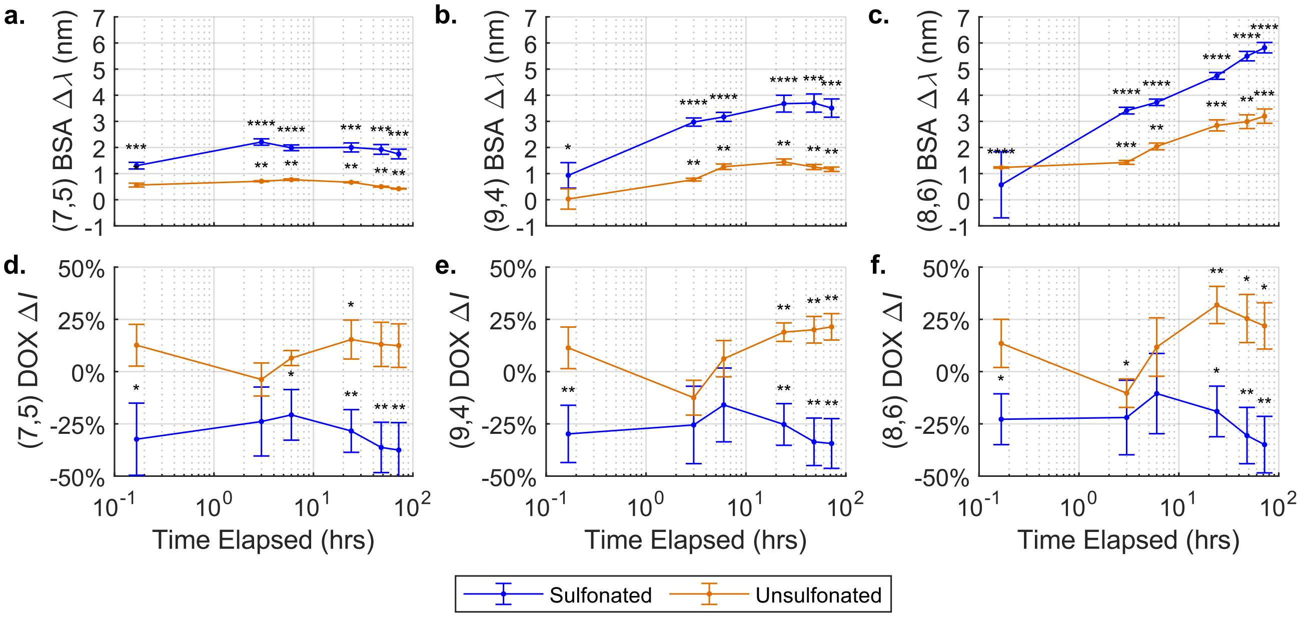


**Figure S4. Effect of Sulfonation on BSA and DOX Detection in MC Gels.** Sulfonated and unsulfonated MC gels were topped with 1X PBS (control), BSA (300 µM) in 1X PBS, or DOX (100 µM) in 1X PBS. Wavelength responses compared at 10 minutes, 3 hours, 6 hours, 24 hours, 48 hours, and 72 hours for a) (7,5) b) (9,4) and c) (8,6) chiralities. Intensity responses compared at same timepoints for a) (7,5) b) (9,4) and c) (8,6) chiralities. N = 4.

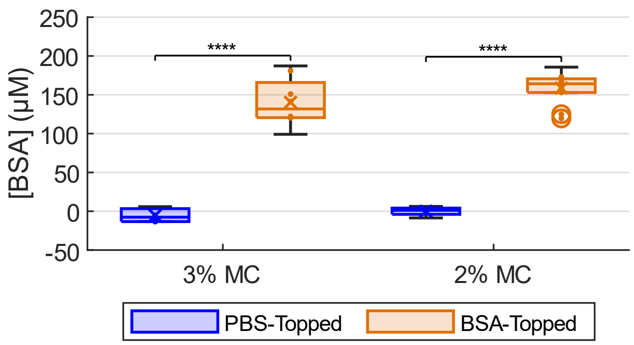


**Figure S5. BSA Content of 2% and 3% MC Gels Post-Topping.** MC gels topped with 1X PBS or BSA (600 µM) in 1X PBS were degraded by cellulase. A BCA assay was then used to quantify the BSA content of the degraded gels, revealing that gels topped with 1X PBS had no BSA content whereas 2% and 3% gels topped with BSA contained 140 ± 30 and 160 ± 20 µM BSA respectively. $p < .0001$

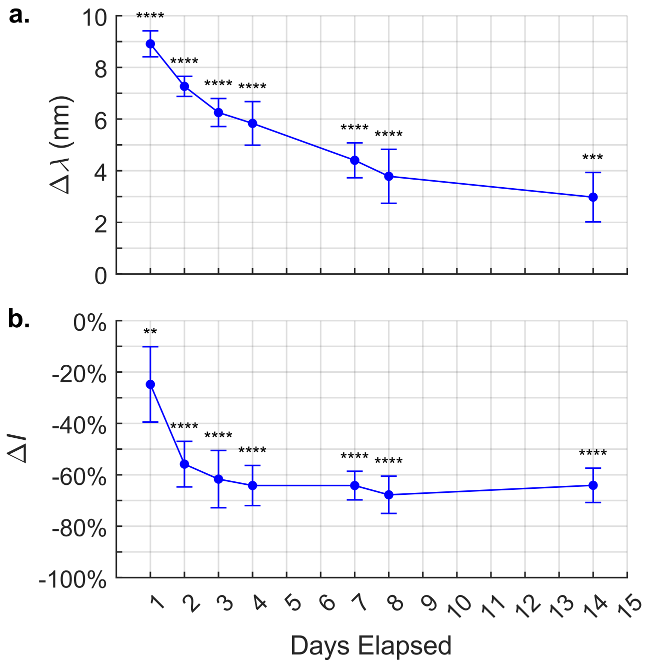


**Figure S6. Long-Term Sensor Response to BSA in 3% Sulfonated MC Hydrogels.** a) Shift (experimental minus control) and b) intensity change (difference of experimental from control as percent of control). All observed wavelength and intensity differences were significant ($p < .01$). Error bars are $\sqrt{variance_{control} + variance_{experimental}}$

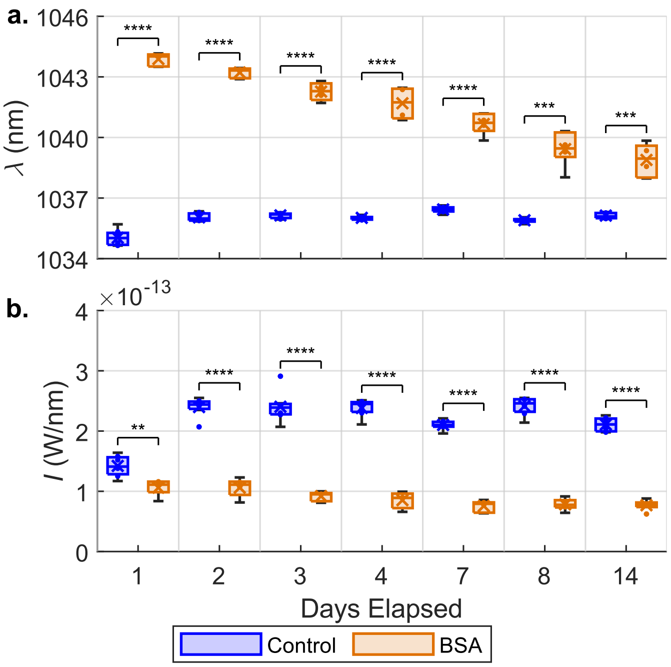


**Figure S7. Response of (GT)$_{15}$-SWCNTs in 3% MC-SO$_3$ to BSA Over Two Weeks.** a) Center wavelength and b) intensity. While all observed wavelength and intensity differences were significant ($p < .01$), highly significant ($p < .0001$) wavelength differences were observed on days 1 through 7, as were intensity differences on days 2-14. ****, ***, **, and * denote $p < .0001$, .001, .01, and .05 respectively.

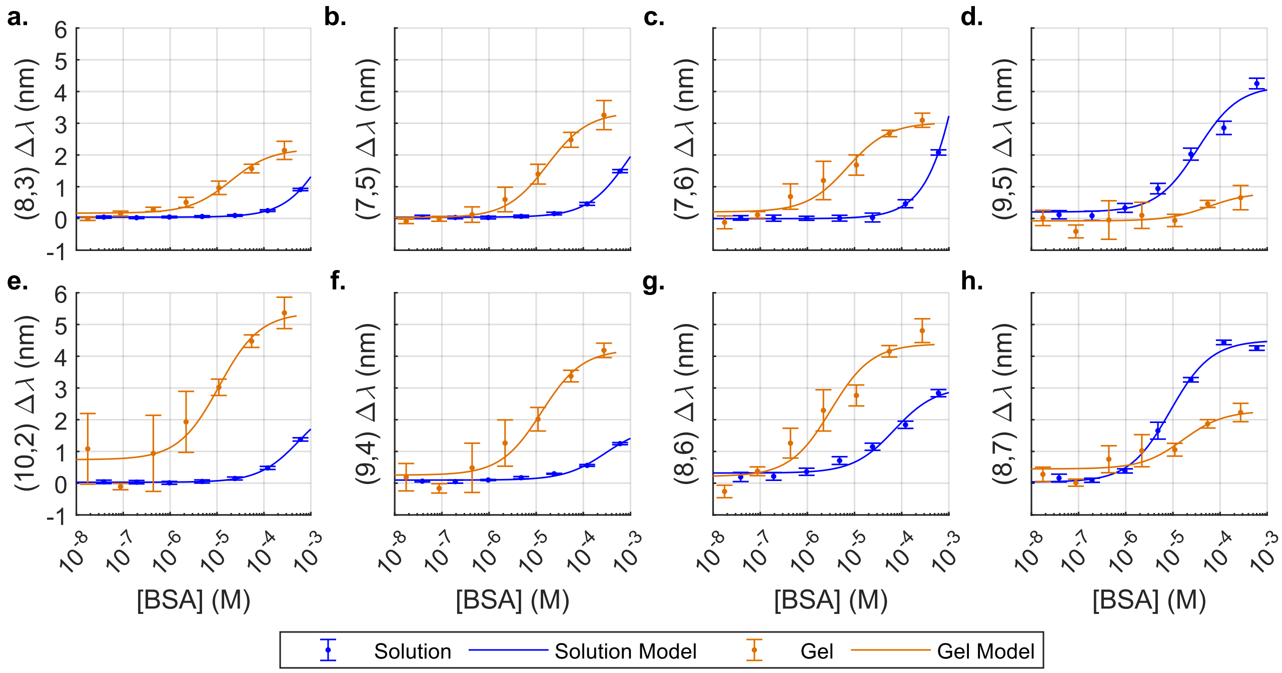


**Figure S8. BSA-Induced Wavelength Response Curves for Various Chiralities of (GT)$_{15}$-SWCNTs in 3% MC-SO$_3$.** a. (8,3) b. (7,5) c. (7,6) d. (9,5) e. (10,2) f. (9,4) g. (8,6) h. (8,7). 0.96 µM gel point excluded from logistic model; gel volume for this group was abnormally high.

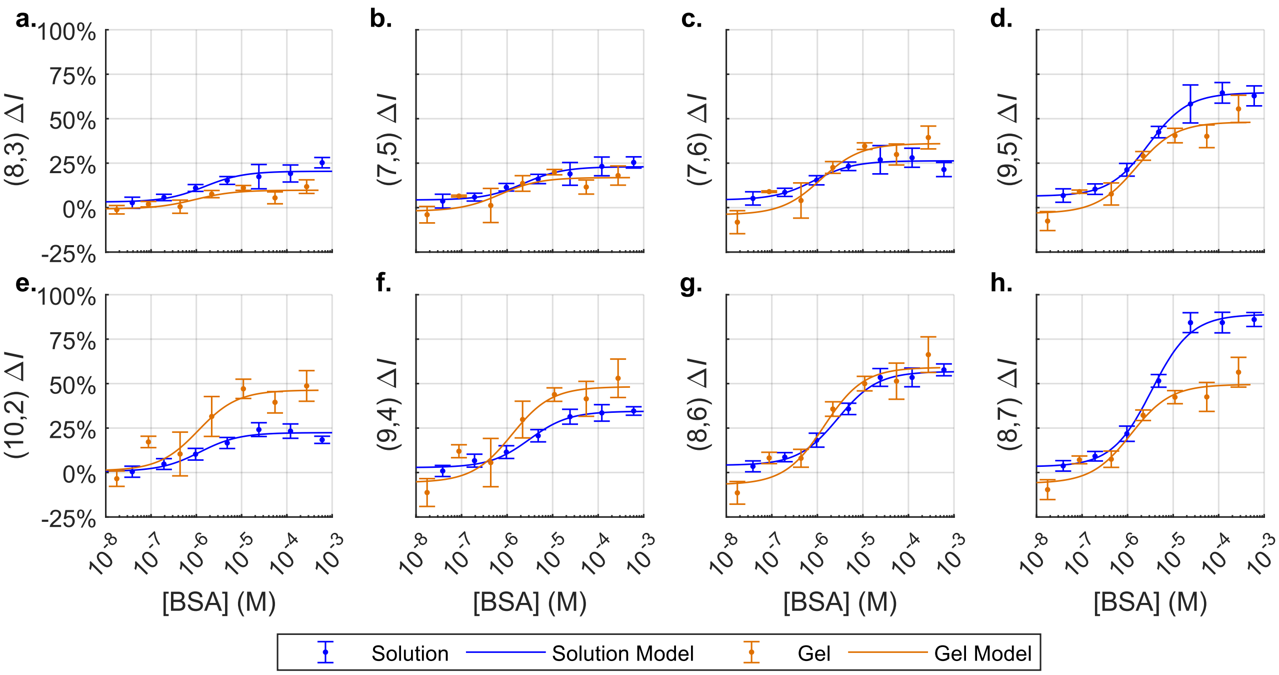


**Figure S9. BSA-Induced Intensity Response Curves for Various Chiralities of $(GT)_{15}$-SWCNTs in 3% MC-$SO_3$.** a. (8,3) b. (7,5) c. (7,6) d. (9,5) e. (10,2) f. (9,4) g. (8,6) h. (8,7). 0.96 µM gel point excluded from logistic model; gel volume for this group was likely abnormally high.

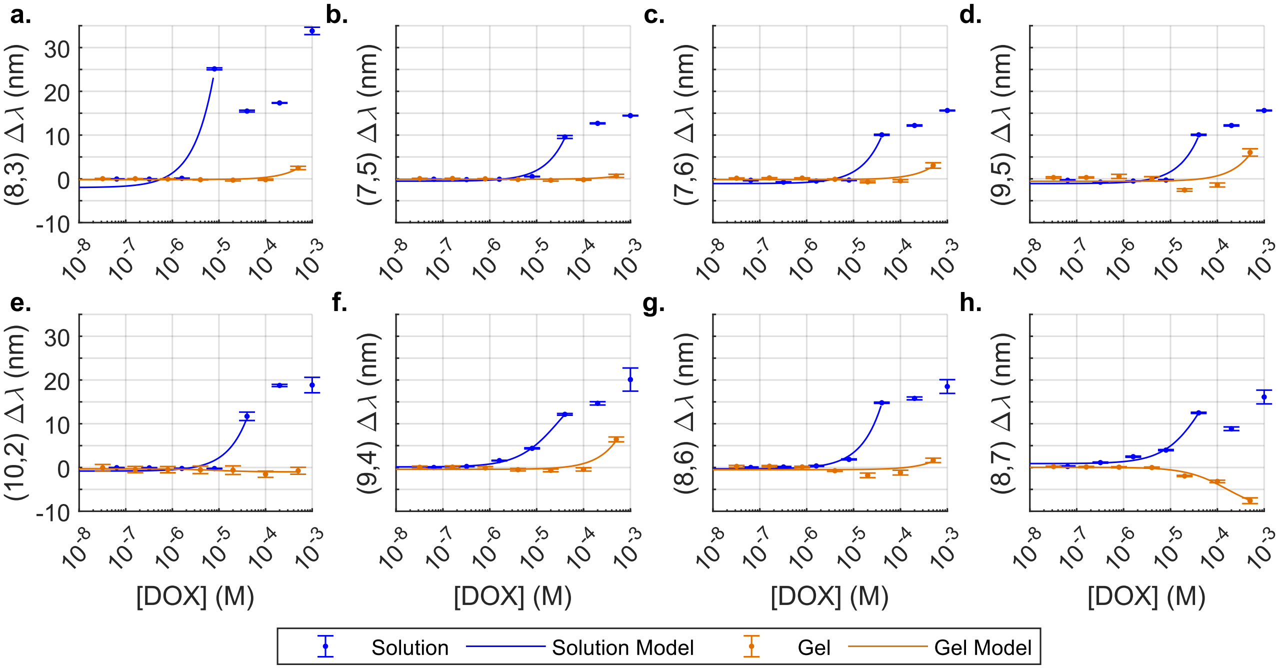


**Figure S10. DOX-Induced Wavelength Response Curves for Chiralities of $(GT)_{15}$-SWCNTs in 3% MC-$SO_3$.** a. (8,3) b. (7,5) c. (7,6) d. (9,5) e. (10,2) f. (9,4) g. (8,6) h. (8,7). For solution data, the model was not extended to concentrations which disobeyed the trend because of DOX aggregation.

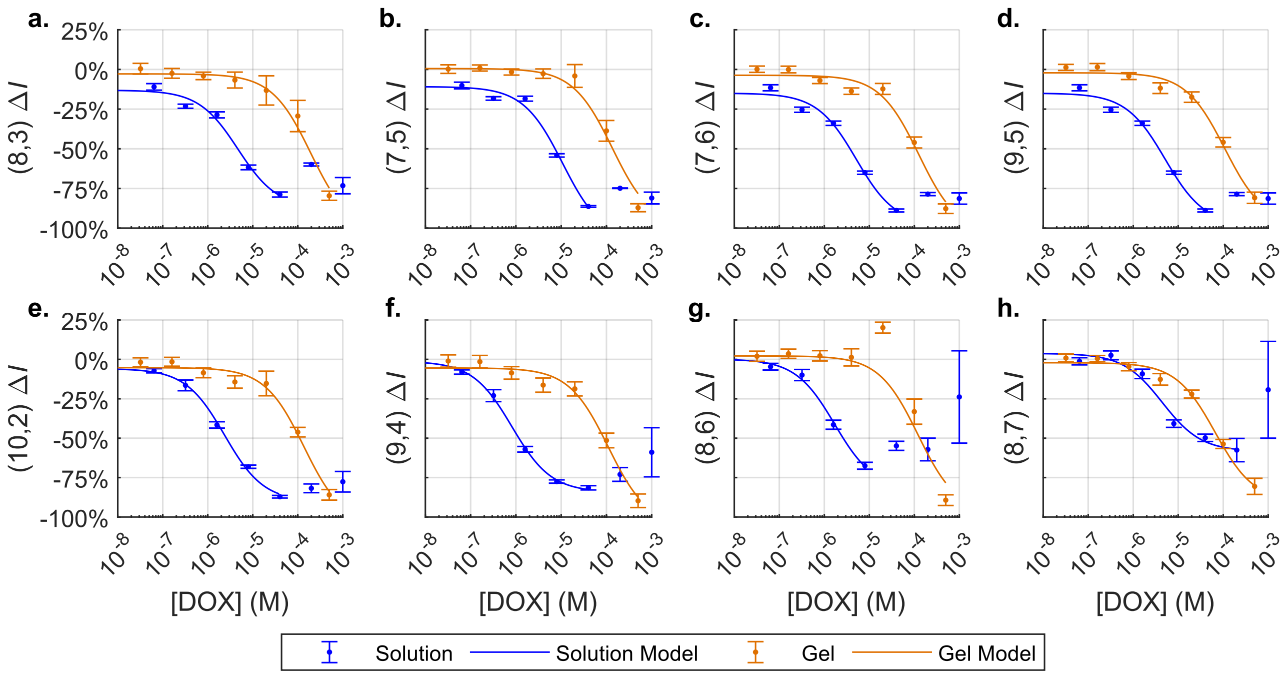


**Figure S11. DOX-Induced Intensity Response Curves for Various Chiralities of (GT)$_{15}$-SWCNTs in 3% MC-SO$_3$.** a. (8,3) b. (7,5) c. (7,6) d. (9,5) e. (10,2) f. (9,4) g. (8,6) h. (8,7). For solution data, the model was not extended to concentrations which disobeyed the trend because of DOX aggregation.

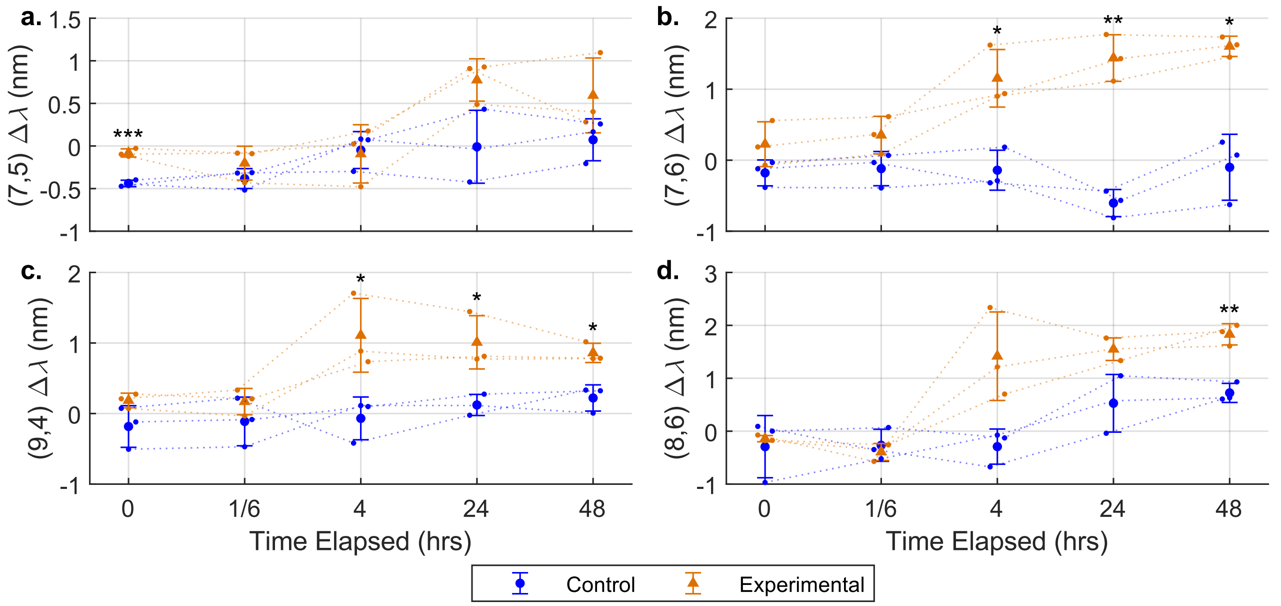


**Figure S12. Shifts of (GT)$_{15}$-SWCNTs in 3% MC-SO$_3$ Implanted in Vivo After Subcutaneous Solution Injection (Experimental and Control).** Error bars show group standard deviation from average. a. (7,5) b. (7,6) c. (9,4) d. (8,6). Shifts calculated vs pre-injection timepoint; significance calculated for experimental shift vs control shift and reported in Table S7.

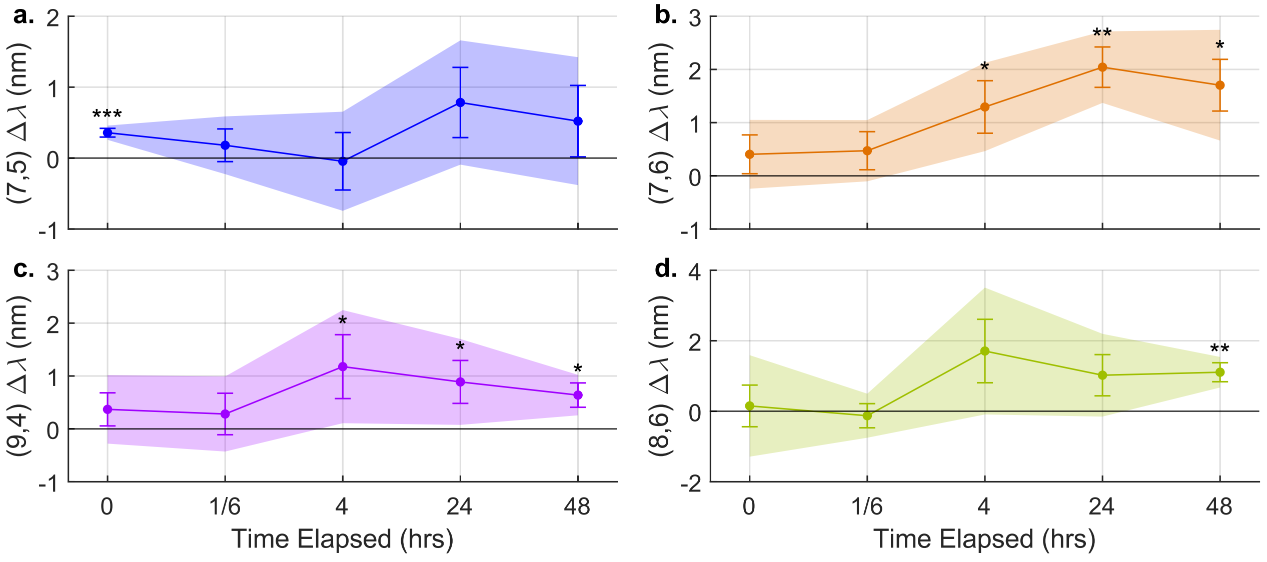


**Figure S13. Shifts of $(GT)_{15}$-SWCNTs in 3% MC-$SO_3$ Implanted in Vivo After Subcutaneous Solution Injection (Difference of Experimental from Control).** Outlines represent 95% confidence interval; error bars represent standard deviation, calculated as $\sqrt{variance_{control} + variance_{experimental}}$ a. (7,5) b. (7,6) c. (9,4) d. (8,6). Shifts calculated vs pre-injection timepoint; significance calculated for experimental shift vs control shift and reported in Table S7.

### Supplementary Tables

**Table S1. Sulfonated Gel Wavelength Response Significance (p value)**

| Days Elapsed | 3% $MgCl_2$ | 3% BSA | 2% $MgCl_2$ | 2% BSA |
|---|---|---|---|---|
| 1 | $2 \times 10^{-6}$ | $5 \times 10^{-6}$ | $5 \times 10^{-5}$ | $4 \times 10^{-5}$ |
| 2 | .003 | .02 | $8 \times 10^{-4}$ | $9 \times 10^{-4}$ |
| 6 | $2 \times 10^{-5}$ | $4 \times 10^{-5}$ | $8 \times 10^{-7}$ | $2 \times 10^{-6}$ |

**Table S2. Sulfonated Gel Intensity Response Significance (p value)**

| Days Elapsed | 3% $MgCl_2$ | 3% BSA | 2% $MgCl_2$ | 2% BSA |
|---|---|---|---|---|
| 1 | $2 \times 10^{-7}$ | $9 \times 10^{-6}$ | $1 \times 10^{-7}$ | $5 \times 10^{-6}$ |
| 2 | | $4 \times 10^{-5}$ | $5 \times 10^{-5}$ | $5 \times 10^{-5}$ |
| 6 | | .02 | | .048 |

**Table S3. Long-term BSA Detection Significance (p value)**

| Days Elapsed | Wavelength | Intensity |
|---|---|---|
| 1 | $1 \times 10^{-14}$ | 0.001 |
| 2 | $3 \times 10^{-13}$ | $1 \times 10^{-9}$ |
| 3 | $2 \times 10^{-8}$ | $2 \times 10^{-6}$ |
| 4 | $1 \times 10^{-6}$ | $7 \times 10^{-11}$ |
| 7 | $9 \times 10^{-7}$ | $1 \times 10^{-12}$ |
| 8 | $6 \times 10^{-5}$ | $4 \times 10^{-10}$ |

| | | |
|---|---|---|
| 14 | $1 \times 10^{-4}$ | $4 \times 10^{-11}$ |

**Table S4. Logistic Model Results: BSA-Induced Wavelength Response.**

| Chirality | Solution | | | Gel | | |
|---|---|---|---|---|---|---|
| | $\Delta_{max}$ (nm) | $K_D$ (μM) | $R^2$ | $\Delta_{max}$ (nm) | $K_D$ (μM) | $R^2$ |
| (8,3) | 4.26 | 2350 | 0.997 | 2.03 | 19.5 | 0.95 |
| (7,5) | 3.67 | 920 | 0.998 | 3.33 | 17.3 | 0.957 |
| (7,6) | 19.5 | 5020 | 0.991 | 2.81 | 7.12 | 0.902 |
| (9,5) | 3.98 | 33.5 | 0.97 | 0.89 | 56.7 | 0.462 |
| (10,2) | 2.65 | 580 | 0.997 | 4.63 | 11.1 | 0.859 |
| (9,4) | 1.70 | 293 | 0.986 | 3.96 | 12.9 | 0.917 |
| (8,6) | 2.71 | 69.6 | 0.975 | 4.18 | 3.15 | 0.902 |
| (8,7) | 4.47 | 8.74 | 0.993 | 1.83 | 16.3 | 0.801 |

**Table S5. Logistic Model Results: BSA-Induced Intensity Response.**

| Chirality | Solution | | | Gel | | |
|---|---|---|---|---|---|---|
| | $\Delta_{max}$ (%) | $K_D$ (μM) | $R^2$ | $\Delta_{max}$ (%) | $K_D$ (μM) | $R^2$ |
| (8,3) | 17.3 | 1.50 | 0.791 | 10.5 | 0.93 | 0.638 |
| (7,5) | 18.7 | 2.33 | 0.839 | 18.9 | 0.70 | 0.652 |
| (7,6) | 22.0 | 0.87 | 0.836 | 40.1 | 1.21 | 0.857 |
| (9,5) | 58.2 | 2.89 | 0.970 | 51.4 | 1.52 | 0.899 |
| (10,2) | 21.8 | 1.20 | 0.895 | 45.4 | 1.11 | 0.831 |
| (9,4) | 31.7 | 3.08 | 0.953 | 53.9 | 1.27 | 0.851 |
| (8,6) | 52.7 | 2.82 | 0.981 | 66.0 | 1.41 | 0.923 |
| (8,7) | 85.6 | 3.36 | 0.987 | 55.6 | 1.20 | 0.916 |

**Table S6. Logistic Model Results: DOX-Induced Intensity Response.**

| Chirality | Solution | | | Gel | | |
|---|---|---|---|---|---|---|
| | $\Delta_{max}$ (%) | $K_D$ (μM) | $R^2$ | $\Delta_{max}$ (%) | $K_D$ (μM) | $R^2$ |
| (8,3) | -74.2 | 4.68 | 0.982 | -100 | 195 | 0.962 |
| (7,5) | -96.3 | 10.7 | 0.986 | -100 | 134 | 0.967 |
| (7,6) | -83.1 | 5.12 | 0.986 | -100 | 126 | 0.976 |
| (9,5) | -83.1 | 5.12 | 0.986 | -95.2 | 108 | 0.983 |
| (10,2) | -84.3 | 2.36 | 0.994 | -100 | 136 | 0.977 |
| (9,4) | -83.1 | 0.82 | 0.996 | -100 | 109 | 0.980 |
| (8,6) | -82.6 | 1.70 | 0.991 | -100 | 121 | 0.966 |
| (8,7) | -62.1 | 4.42 | 0.971 | -88.2 | 66.0 | 0.985 |

**Table S7. Significance of DOX-Induced Shifting in vivo (p value)**

| Hours Elapsed | (7,5) | (7,6) | (9,4) | (8,6) |
|---|---|---|---|---|
| 0 | .007 | N.S. | N.S. | N.S. |
| 4 | N.S. | .01 | .04 | N.S. |
| 24 | N.S. | .002 | .04 | N.S. |
| 48 | N.S. | .02 | .01 | .002 |

#### Supplementary Equations

$$I_{\lambda} = I_{max}\left(\frac{1}{1+\left(\frac{\lambda-\lambda_0}{\frac{\gamma}{2}}\right)^2} \times e^{-\ln(2)\left(\frac{\lambda-\lambda_0}{\frac{\sigma}{2}}\right)^2}\right) + m\lambda + b$$

(Equation S1)

Where

$I_{\lambda}$ is the fluorescence intensity at wavelength $\lambda$

$I_{max}$ is the baseline-subtracted fluorescence intensity at the center wavelength

$\lambda_0$ is the center wavelength

$\gamma$ is the full-width-at-half-maximum of the Lorentzian component

$\sigma$ is the full-width-at-half-maximum of the Gaussian component

$m$ is the slope of the baseline

$b$ is the y-intercept of the baseline

$$response = baseline + \frac{\Delta_{max} \times [analyte]}{K_D + [analyte]}$$

(Equation S2)

Where

*response* is the change in the signal (center wavelength or intensity) at a given analyte concentration

*baseline* is the response of the control.

$\Delta_{max}$ is the maximum response

$K_D$ is the analyte concentration which elicits 50% of the maximum response. Assuming the sensor response is directly determined by the amount of bound analyte, this should be equal to the dissociation constant.

*[analyte]* is the analyte concentration

#### References

(1) Gold, G. T.; Varma, D. M.; Taub, P. J.; Nicoll, S. B. Development of crosslinked methylcellulose hydrogels for soft tissue augmentation using an ammonium persulfate-ascorbic acid redox system. *Carbohydr Polym* **2015**, *134*, 497-507. DOI: 10.1016/j.carbpol.2015.07.101  From NLM Medline.
